# Bayesian inference in ring attractor networks

**DOI:** 10.1101/2021.12.17.473253

**Authors:** Anna Kutschireiter, Melanie A Basnak, Jan Drugowitsch

**Affiliations:** Harvard Medical School, Department of Neurobiology, 200 Longwood Avenue, Boston, MA- 02115, United States

**Keywords:** Working memory, Bayesian inference, Ring attractor networks, Head direction neurons, Kalman filter, Population coding, Bump attractor

## Abstract

Working memories are thought to be held in attractor networks in the brain. These attractors should keep track of the uncertainty associated with each memory, so as to weigh it properly against conflicting new evidence. However, conventional attractors do not represent uncertainty. Here we show how uncertainty could be incorporated into an attractor, specifically a ring attractor that encodes head direction. First, we introduce the first rigorous normative framework (the circular Kalman filter) for benchmarking the performance of a ring attractor under conditions of uncertainty. Next we show that the recurrent connections within a conventional ring attractor can be re-tuned to match this benchmark. This allows the amplitude of network activity to grow in response to confirmatory evidence, while shrinking in response to poor-quality or strongly conflicting evidence. This “Bayesian ring attractor” performs near-optimal angular path integration and evidence accumulation. Indeed, we show that a Bayesian ring attractor is consistently more accurate than a conventional ring attractor. Moreover, near-optimal performance can be achieved without exact tuning of the network connections. Finally, we use large-scale connectome data to show that the network can achieve near-optimal performance even after we incorporate biological constraints. Our work demonstrates how attractors can implement a dynamic Bayesian inference algorithm in a biologically plausible manner, and it makes testable predictions with direct relevance to the head direction system, as well as any neural system that tracks direction, orientation, or periodic rhythms.

**Significance Statement:** Data from human subjects as well as animals shows that working memories are associated with a sense of uncertainty. Indeed, a sense of uncertainty is what allows an observer to properly weigh new evidence against their current memory. However, we do not understand how the brain tracks uncertainty. Here we describe a simple and biologically plausible network model that can track the uncertainty associated with a working memory. The representation of uncertainty in this model improves the accuracy of its working memory, as compared to conventional models, because it assigns the proper weight to new conflicting evidence. Our model provides a new interpretation for observed fluctuations in brain activity, and it makes testable new predictions.

## Introduction

Attractor networks are thought to form the basis of working memory (Wang 2001; Compte 2006) as they can exhibit persistent, stable activity patterns (attractor states) even after network inputs have ceased (Hansel and Sompolinsky 1998). An attractor network can gravitate toward a stable state even if its input is based on partial (unreliable) information; this is why attractors have been suggested as a mechanism for pattern completion (Hopfield 1982). However, the characteristic stability of any attractor network also creates a problem: once the network has settled into its attractor state, it will no longer be possible to see that its inputs might have been unreliable. In this situation, the attractor state will simply represent a point estimate (or “best guess”) of the remembered input, without any associated sense of uncertainty. However, real memories often include a sense of uncertainty (e.g., (Rademaker et al. 2012; Li et al. 2021)), and uncertainty has clear behavioral effects (Ernst and Banks 2002; Piet et al. 2018; Fetsch et al. 2009). This motivates us to ask how an attractor network can conjunctively encode a memory and its associated uncertainty.

A ring attractor is a special case of an attractor that can encode a circular variable (Knierim and Zhang 2012). For example, there is good evidence that the neural networks that encode head direction (HD) are ring attractors (Zhang 1996; Skaggs et al. 1994; Redish et al. 1996; Peyrache et al. 2015; Seelig and Jayaraman 2015; Kim et al. 2017; Turner-Evans et al. 2017; Ajabi et al. 2021). In a conventional ring attractor, inputs push a “bump” of activity around the ring, with only short-lived changes in bump amplitude or shape (Amari 1977; Ermentrout 1998); the rapid decay to a stereotyped bump shape is by design, such that this type of conventional ring attractor network is unable to track uncertainty. However, it would be useful to modify these conventional ring attractors so that they can encode the uncertainty associated with HD estimates. HD estimates are constructed from two types of observations — angular velocity observations and HD observations (Knierim and Zhang 2012; Heinze et al. 2018). Angular velocity observations arise from vestibular or proprioceptive signals, as well as optic flow; these observations indicate the head’s rotational movement, and thus a change in HD (Skaggs et al. 1994; Xie et al. 2002; Turner-Evans et al. 2017; Green et al. 2017). These angular velocity observations are integrated over time (“remembered”) to update the system’s internal estimate of HD, in a process termed angular path integration. Ideally, a ring attractor would track the uncertainty associated with angular path integration errors. Meanwhile, HD observations arise from visual landmarks or other sensory cues that provide information about the head’s current orientation (Zhang 1996; Seelig and Jayaraman 2015). These sensory observations can change the system’s internal HD estimate, and once that change has occurred, it is generally persistent (remembered). But like any sensory signal, these sensory observations are noisy; they are not unambiguous evidence of HD. Therefore, the way that a ring attractor responds to each new visual landmark observation should ideally depend on the uncertainty associated with its current HD estimate. This type of uncertainty-weighted cue integration is a hallmark of Bayesian inference (Knill and Pouget 2004), and would require a network to keep track of its own uncertainty.

In this study, we address three related questions. First, how should an ideal observer integrate uncertain evidence over time to estimate a circular variable? For a linear variable, this is typically done with a Kalman filter; here we introduce an extension of Kalman filtering for circular statistics; we call this the circular Kalman filter. This algorithm provides a high-level description of how the brain *should* integrate evidence over time to estimate HD, or indeed any other circular or periodic variable. Second, how could a neural network actually implement the circular Kalman filter? We show how this algorithm could be implemented by a neural network whose basic connectivity pattern resembles that of a conventional ring attractor. With properly tuned network connections, we show that the bump amplitude grows in response to confirmatory evidence, whereas it shrinks in response to strongly conflicting evidence, or the absence of evidence. We call this network a Bayesian ring attractor. Third, how does the performance of a Bayesian ring attractor compare to the performance of a conventional ring attractor? In a conventional ring attractor, bump amplitude is pulled rapidly back to a stable baseline value, whereas in a Bayesian ring attractor, bump amplitude is allowed to float up or down as the system’s certainty fluctuates. As a result, we show that a Bayesian ring attractor has consistently more accurate internal estimates (or “working memory”) of the variable it is designed to encode than a conventional ring attractor.

Together, these results provide a principled theoretical foundation for how ring attractor networks can be tuned to conjointly encode a memory and its associated uncertainty. Although we focus on the brain’s HD system as a concrete example, our results are relevant to any other brain system that encodes a circular or periodic variable.

## Results

### Circular Kalman filtering: a Bayesian algorithm for tracking a circular variable

We begin by asking how an ideal observer should dynamically integrate uncertain evidence to estimate a circular variable. At each time point, the observer’s estimate is represented by a probability distribution on a circle, where the peak of the distribution represents the best guess, and the width of the distribution represents the uncertainty associated with that guess (Fig. 1a). This distribution is termed a posterior belief (Knill and Pouget 2004; Dehaene et al. 2021). Each new observation updates this belief. Importantly, any given observation has limited reliability, that is, it provides incomplete information. Therefore, we also characterize these observations by probability distributions. In the brain’s HD system, an estimate of head direction (*ϕ*) is based on angular velocity observations (*ν*) as well as HD observations (*z*), each with associated reliability (*κ_ν_* and *κ_z_*). What limits reliability is noise; specifically, we assume that angular velocity observations are corrupted by Gaussian noise, while direction observations are corrupted by von Mises noise (because a directional variable is circular, and von Mises approximates Gaussian on a circle). The distribution of probable head directions *p*(*ϕ_t_*|*z*_0:*t*_, *ν*_0:*t*_) at current time *t* depends on the accumulated history of these observations up to time *t*.

**Figure 1.**
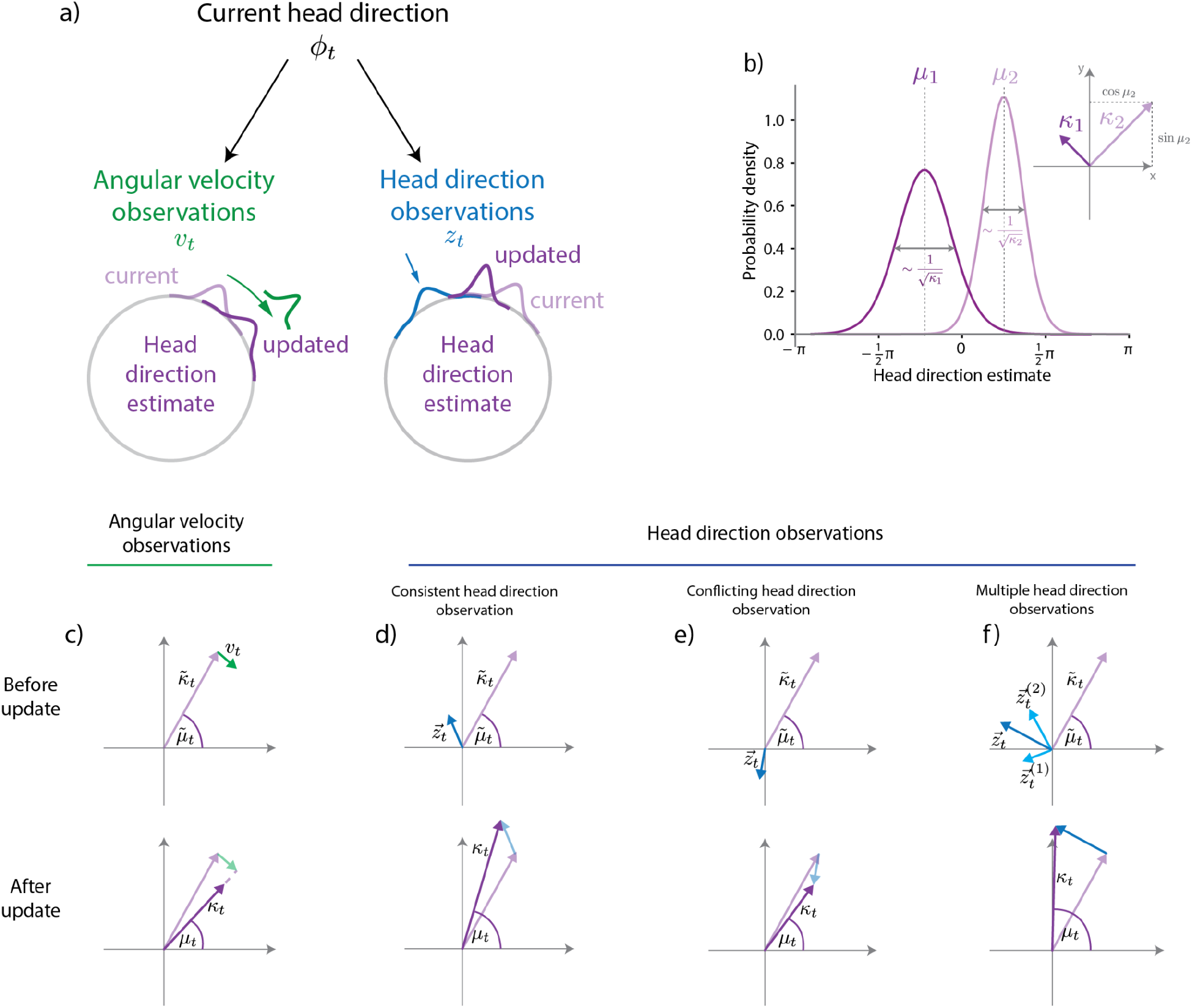
Tracking HD with the circular Kalman filter. **a)** Angular velocity observations provide noisy information about the true angular velocity 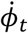, while HD observations provide noisy information about the true HD *ϕ*_t_. **b)** At every point in time, the posterior belief *p*(*ϕ_t_*) is approximated by a von Mises distribution. It is fully characterized by its mean parameter *μ_t_*, which determines the position of the peak, and its certainty parameter *κ_t_*. Interpreted as the polar coordinates in the 2D plane, these parameters provide a convenient vector representation of the posterior belief (inset). **c)** A angular velocity observation *ν_t_* is a vector tangent to the current HD estimate. Angular velocity observations continually rotate the current HD estimate; meanwhile, noise accumulation progressively decreases certainty. **d)** Each HD observation *z_t_* is a vector whose length corresponds to the reliability associated with that observation. Adding this vector to the current HD estimate produces an updated HD estimate. If the HD observation is compatible with the current HD estimate, then the updated HD estimate has increased certainty. **e)** HD observations in conflict with the current belief (e.g., opposite direction of current estimate) decrease the certainty associated with the HD estimate. **f)** Multiple HD cues can be integrated simultaneously via vector addition.

The process of updating an estimate based on noisy observations is often accomplished using a Kalman filter (Kalman 1960; Kalman and Bucy 1961), but this assumes that the encoded variable is linear; a circular variable requires a different approach. Because filtering on a circle is analytically intractable (Kurz et al. 2016), we choose to approximate the distribution of probable head directions by a von Mises distribution, with mean *μ* and certainty (i.e., precision) *κ*, so that *P*(*ϕ_t_*|*z*_0:*t*_, *ν*_0:*t*_) ≈ *VM*(*ϕ_t_*|*μ_t_*, *κ_t_*). This allows us to update the estimated circular variable **ϕ*_t_* using a technique called projection filtering (Brigo et al. 1999; Kutschireiter et al. 2022):

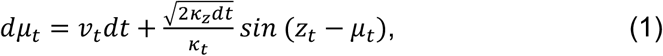

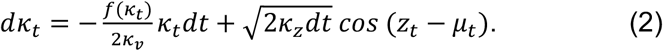

Here, *f*(*κ_t_*) is a monotonically increasing nonlinear function that controls the speed of decay in certainty *κ_t_* (see Methods). Equations (1) and (2) describe an algorithm that we call the **circular Kalman filter** (Kutschireiter et al. 2022). This algorithm provides a general solution for estimating the evolution of a circular variable over time, based on noisy observations.

To understand the circular Kalman filter intuitively, it is helpful to think of the observer’s estimate as a vector (Fig. 1b), whose direction represents the current best guess μ_t_, and whose length represents the certainty *κ_t_* associated with that guess. The circular Kalman filter tells us how this vector should change at each time point, based on new observations of angular velocity and direction. Here we outline the intuition behind the circular Kalman filter, focusing on the HD system as a specific example.

#### Angular velocity observations

We can think of each angular velocity observation as a vector which points at a tangent to the current HD estimate (Fig. 1c) and rotates the current HD estimate (first RHS term in Eq. (1)). Angular velocity observations are noisy and so decrease the HD estimate’s certainty (*κ_t_*), meaning that the observer’s estimate vector becomes shorter (Fig. 1c). Thus, when angular velocity observations are the only inputs to the HD network — i.e., when HD observations are absent — the certainty *κ_t_* associated with the HD estimate will progressively decay, with a speed of decay that depends on *κ_ν_* (first RHS term in Eq. (2)).

#### HD observations

We can treat each HD observation as a vector, with a length *κ_z_* representing the reliability associated with that observation (e.g., the reliability of a visual landmark observation). We add the HD observation vector to the current HD estimate to obtain the updated HD estimate. The updated estimate’s direction depends on the relative lengths of both vectors. A relatively longer HD observation vector, i.e., a more reliable observation relative to the current HD estimate’s certainty, results in a stronger impact on the updated HD estimate (Fig. 1d, second RHS term in Eq. (1)). In line with principles of reliability-weighted Bayesian cue combination (Knill and Pouget 2004) HD observations increase the observer’s certainty if they are confirmatory (i.e., they indicate that the current estimate is correct or nearly so, Fig. 1d). Interestingly, however, if HD observations strongly conflict with the current estimate, they actually *decrease* certainty (Fig. 1e). This notable result is a consequence of the circular nature of the inference task (Murray and Morgenstern 2010). It stands in contrast to the standard (non-circular) Kalman filter, where an analogous observation would *always* increase the observer’s certainty (Wilson and Finkel 2009). It is thus a key distinction between the standard Kalman filter and the circular Kalman filter.

To summarize, the circular Kalman filter describes how an ideal observer should integrate a stream of unreliable information over time to update an estimate of a circular variable. This algorithm serves as a normative standard to judge the performance of any network in the brain that tracks a circular or periodic variable. Specifically, in the HD system, the circular Kalman filter tells us that angular velocity observations should rotate the HD estimate while reducing the certainty of that estimate. Meanwhile, HD observations should update the HD estimate weighted by their reliability, and they should either increase certainty (if compatible with the current estimate) or reduce it (if strongly conflicting with the current estimate). Note that the circular Kalman filter can integrate HD observations from multiple sources by simply adding all their vectors to the current HD estimate vector (Fig. 1f).

### Neural encoding of a probability distribution

Thus far, we have developed a normative algorithmic description of how an observer *should* integrate evidence over time to estimate a circular variable. This algorithm requires the observer to represent their current estimate as a probability distribution on a circle. How could a neural network encode this probability distribution? Consider a ring attractor network where adjacent neurons have adjacent tuning preferences, so that the population activity pattern is a spatially localized “bump”. The bump’s center of mass is generally interpreted as a point estimate (or best guess) of the encoded circular variable (Ben-Yishai et al. 1995; Zhang 1996). In the HD system, this would be the best guess of head direction. Meanwhile, we let the bump amplitude encode certainty, so that higher amplitude corresponds to higher certainty. Of course, there are other ways to encode certainty — e.g., using bump width rather than bump amplitude. However, there are two good reasons for focusing on bump amplitude. First, as we will see below, this implementation allows the parameters of the encoded probability distribution to be “read out” in a particularly straightforward way. Second, recent data from the mouse HD system shows that the appearance of a visual cue (which increases certainty) causes bump amplitude to increase; moreover, when the bump amplitude is high, the network is relatively insensitive to the appearance of a visual cue that conflicts with the current HD estimate, again suggesting that bump amplitude is a proxy for certainty (Johnson et al. 2005; Ajabi et al. 2021).

Formally, then, the activity of a neuron i with preferred HD *ϕ_i_* can be written as follows (Fig. 2a):

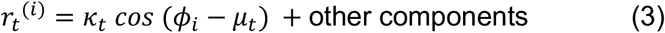

where *μ_t_* is the HD point estimate, *κ_t_* is the associated certainty, and the “other components” might include a constant (representing baseline activity) or minor contributions of higher-order Fourier components. Note that Eq. (3) does not imply that the tuning curve must be cosine-shaped. Rather, it implies that the cosine *component* of the tuning curve is scaled by certainty. This is satisfied, for example, by any unimodal bump profile whose overall gain is governed by certainty. A particularly interesting case that matches Eq. (3) is a linear probabilistic population code (Ma et al. 2006; Beck et al. 2011) with von Mises-shaped tuning curves and independent Poisson neural noise (SI and Fig. S1).

**Figure 2.**
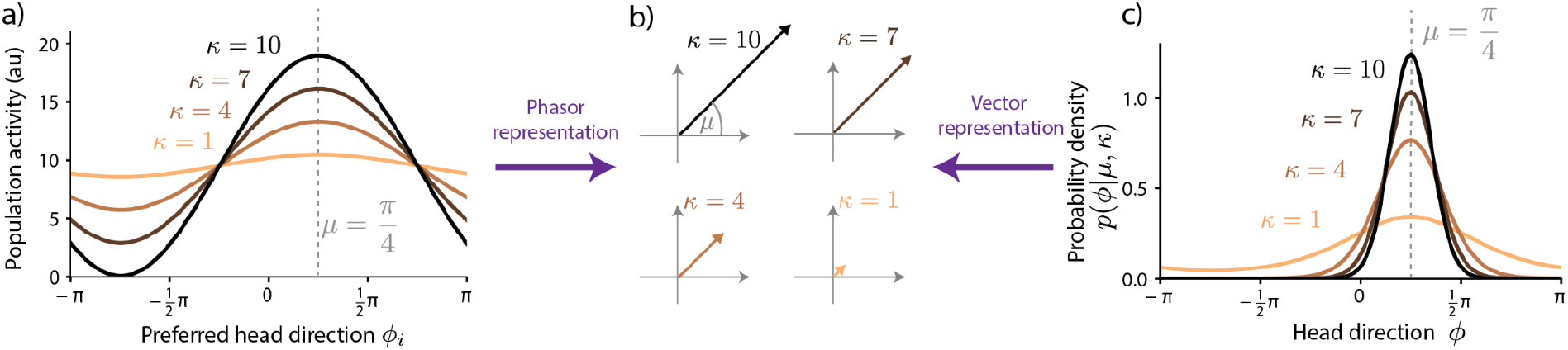
Encoding the HD belief in neural population activity. **a)** Neural population activity profile (e.g., average firing rate) encoding the HD estimate *μ* = *π*/4 with different values of certainty κ. Neurons are sorted by preferred head directions *ϕ_i_*. **b)** Vector representation of estimate *μ* = *π*/4 for different values of certainty *κ*. This vector representation can be obtained by linearly decoding the population activity in a) (“phasor representation”). It also corresponds to the vector representation of the von Mises distribution in c). Thus, it links neural activities to the probability distributions they encode. **c)** Von Mises probability densities for different values of certainty *κ* and fixed HD estimate *μ* = *π*/4. Note that, unlike the population activity in a), the density sharpens around the mean with increasing certainty.

Importantly, this neural representation would allow downstream neurons to read out the parameters of the probability distribution *p*(*ϕ_t_*|*z*_0:*t*_, *ν*_0:*t*_) in a straightforward manner. Specifically, downstream neurons could take a weighted sum of the population firing rates (Methods) to recover two parameters, *θ*_1_ = *κ_t_ cos* (*μ_t_*) and *θ*_2_ = *κ_t_ sin* (*μ_t_*). In other words, the parameters *θ_1_* and *θ*_2_ would be accessible to downstream neurons via simple (linear) neural operations. This is notable because *θ*_1_ and *θ*_2_ represent the von Mises distribution *p*(*ϕ_t_*|*z*_0:*t*_, *ν*_0:*t*_) in terms of Cartesian vector coordinates in the 2D plane, whereas μ and *κ* are its polar coordinates (Fig. 1b). This is related to the phasor representation of neural activity (Lyu et al. 2022), which also translates bump position and amplitude to polar coordinates in the 2D plane (Fig. 2b). If the amplitude of the activity bump scales with certainty, the phasor representation of neural activity equals the vector representation of the von Mises distribution (Fig 2b,c).

### Neural network implementation of the circular Kalman filter

Now that we have specified how our model network represents the probability distribution *p*(*ϕ_t_*|*z*_0:*t*_, *ν*_0:*t*_) over possible head directions, we can proceed to considering the dynamics of this network — specifically, how it responds to incoming information, or the lack of information. The circular Kalman filter algorithm describes the vector operations required to dynamically update the probability distribution *p*(*ϕ_t_*|*z*_0:*t*_, *ν*_0:*t*_, *ν*_0:*t*_) with each new observation of angular velocity or head direction. Here we show how the circular Kalman filter could be implemented by a neural network, which we call a Bayesian ring attractor. We describe the features of this network with regard to the HD system, but the underlying concepts are general ones which could be applied to any network that encodes a circular or periodic variable. The dynamics of the Bayesian ring attractor network are given by

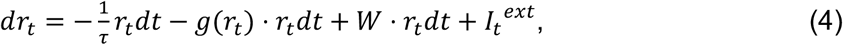

where *r_t_* denotes a population activity vector, with neurons ordered by their preferred head directions *ϕ_i_*, *τ* is the cell-intrinsic leak time constant, *D* is the matrix of excitatory recurrent connectivity, *I_t_^ext^* is a vector of HD observations, and *g* is a nonlinear function that determines global inhibition, and that we discuss in more detail further below. Let us now consider each of these terms in detail.

First, HD observations enter the network via the input vector *I_t_^ext^*, in the form of a cosine-shaped spatial pattern whose amplitude scales with reliability *κ_z_* (Fig. 3a). This implements the vector addition required for the proper integration of these observations. Specifically, the weight assigned to each HD observation is determined by the amplitude of *I_t_^ext^*, relative to the amplitude of the activity bump in the HD population. Thus, observations are weighted by their reliability, relative to the certainty of the current HD estimate, as in the circular Kalman filter (Fig. 1d-e). An HD observation that tends to confirm the current HD estimate will increase the amplitude of the bump in HD cells, and thus the estimate’s certainty.

**Figure 3.**
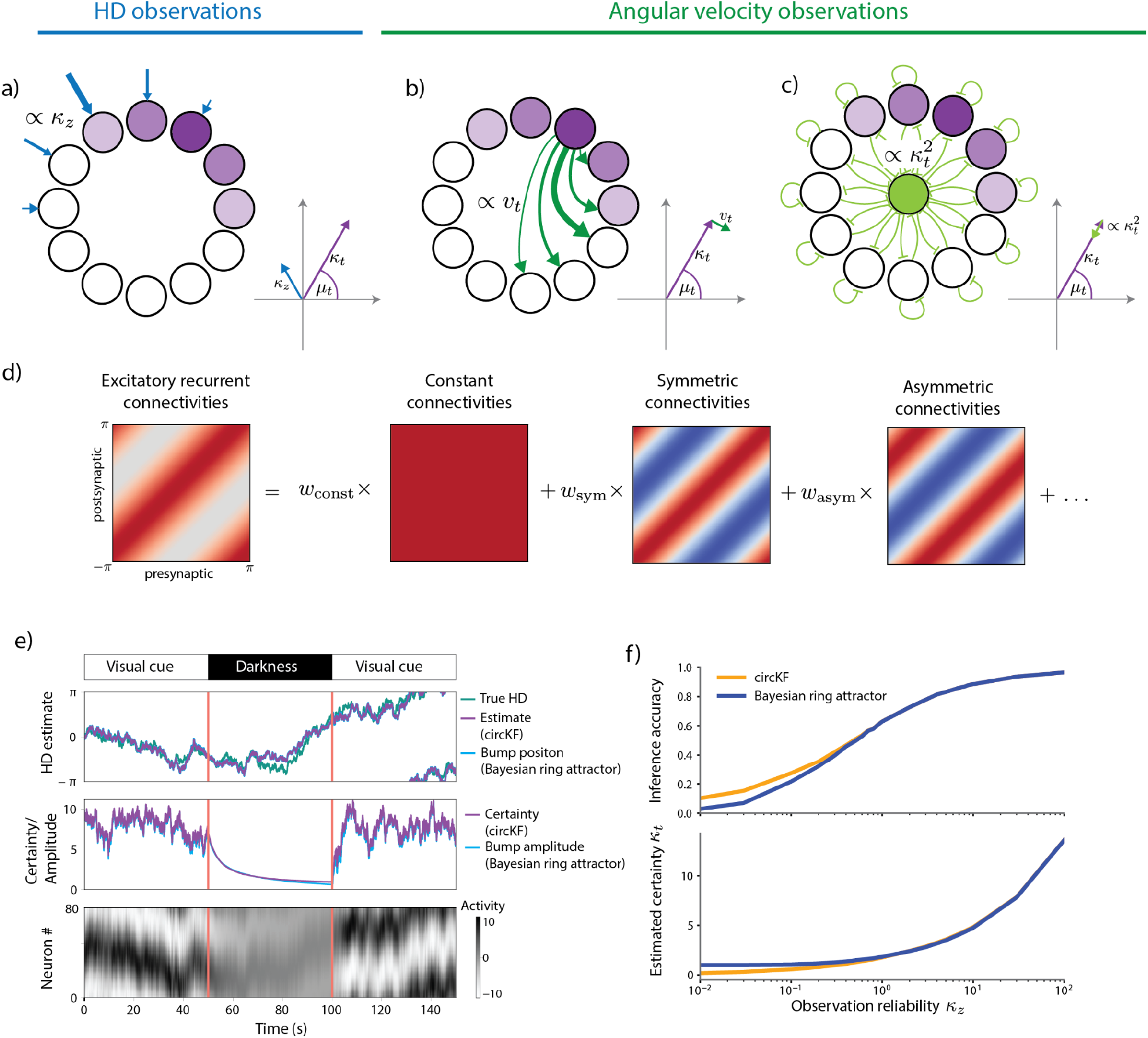
A recurrent neural network implementation of the circular Kalman filter. **a)-c):** Network motifs sufficient to implement the circular Kalman filter. **a**) A cosine-shaped input to the network provides HD observation input. The strengths of this input is modulated by observation reliability *κ_z_*. **b)** Rotations of the HD estimate are mediated by symmetric recurrent connectivities, whose strength is modulated by angular velocity observations. **c)** Decay in amplitude, which implements decreasing certainty in HD estimate, arises from leak and global inhibition. **d)** Rotation-symmetric recurrent connectivities (here: neurons are sorted according to their preferred HD) can be decomposed into constant, symmetric, asymmetric, and higher-order frequency components (here denoted by dots). Red and blue denote excitatory and inhibitory components, respectively. **e)** The dynamics of the Bayesian ring attractor implement the dynamics of the ideal observer’s belief, as shown in a simulation of a network with 80 neurons. Here, we assume that vision provides the network with HD observations. When a ‘visual cue’ was present, both HD observations and angular velocity observations were available. During ‘darkness’, only angular velocity observations were available. **f)** The Bayesian ring attractor network tracks the HD estimate with the same precision (top; higher = lower circular distance to true HD) as the circular Kalman filter (circKF, Eqs. (1) and (2)) if HD observations are reliable (large *κ_z_*), but with slightly lower precision once they become less reliable (small *κ_z_*). This drop co-occurs with an overestimate in the estimate’s confidence *κ_t_* (bottom). The shown accuracies and certainties are averages over 5,000 simulation runs (see Methods for simulation details).

Second, the matrix of recurrent connectivity *W* has spatially symmetric and asymmetric components (Fig. 3d). The symmetric component consists of local excitatory connections that each neuron makes onto adjacent neurons with similar HD preferences. This holds the bump of activity at its current location in the absence of any other input. The overall strength of the symmetric component (*w_sym_*) is a free parameter which we can tune. Meanwhile, the asymmetric component consists of excitatory connections that each neuron makes onto adjacent neurons with shifted HD preferences. This component tends to push the bump of activity around the ring (Fig. 3b). Angular velocity observations modulate the overall strength of the asymmetric component (*w_asym_*), so that positive and negative angular velocity observations push the bump in opposite directions.

Third, decreasing certainty in the HD estimate arising from an increasing angular path integration error is implemented by the global inhibition term, −*g*(*r_t_*) · *r_t_* (Fig. 3c). Here, the function *g*’s output increases linearly with bump amplitude in the HD population, resulting in an overall quadratic inhibition (see Methods). Together with the leak, this quadratic inhibition approximates the nonlinear certainty decay *f*(*κ_t_*) in the circular Kalman filter (Eq. (2)). The approximation becomes precise in the limit of large posterior certainties *κ_t_*.

With the appropriate parameter values, the amplitude of the bump decays slowly as long as new HD observations are unavailable, because global inhibition and leak work together to pull the bump amplitude slowly downward (Fig. 3e). This is by design: the circular Kalman filter tells us that certainty decays slowly without a continuous stream of new HD observations. This situation differs from conventional ring attractors, whose bump amplitudes are commonly designed to rapidly decay to their stable (attractor) states. In a hypothetical network that perfectly implemented the circular Kalman filter, the bump amplitude would decay to zero. However, in our Bayesian ring attractor, which merely approximates the circular Kalman filter, the bump amplitude decays to a low but nonzero baseline amplitude (*κ*\*).

As an illustrative example, we simulated a network of 80 HD neurons (see Methods). We let HD follow a random walk (diffusion on a circle), and we used the time derivative of HD (angular velocity) to modulate the asymmetric component of the connectivity matrix *W*. As HD changes, we rotate the cosine-shaped bump in the external input vector *I_t_^ext^*, simulating the effect of a visual cue whose position on the retina depends on HD. This network exhibits a spatially localized bump whose position tracks HD, with an accuracy similar to that of the circular Kalman filter itself (Fig. 3e). Meanwhile, the amplitude of the bump accurately tracks the fluctuating certainty of the HD estimate in the circular Kalman filter, reflecting how noisy angular velocity and HD observations interact to modulate this certainty (see Eq. (2)). When the visual cue is removed, the bump amplitude decays toward baseline (Fig. 3e). In the limit of infinitely many neurons, this type of network can be tuned to implement the circular Kalman filter exactly for sufficiently high HD certainties. What this simulation shows is that network performance can come close to benchmark performance even with a relatively small number of neurons (Fig. S2).

Interestingly, when we vary the certainty associated with HD observations, we can observe two operating regimes in the network. When HD observations have high certainty (high *κ_z_*), bump amplitude is high and accurately tracks changes HD certainty (*κ_z_*). Thus, in this regime, the network performs proper Bayesian inference (Fig. 3f). Conversely, when HD observations have low certainty (low *κ_z_*), bump amplitude is low but constant, because it is essentially pegged to its baseline value (the network’s attractor state). In this regime, bump amplitude exaggerates the certainty of the HD estimate, and the network looks more like a conventional ring attractor. We will analyze these two regimes further in the next section.

### Bayesian versus conventional ring attractors

Conventional ring attractors (Zhang 1996; Compte 2000; Wang 2001) are commonly designed to operate close to their attractor states, so that bump amplitude is nearly constant. This is not true of the Bayesian ring attractor described above, where bump amplitude varies by design. The motivation for this design choice was the idea that, if bump amplitude varies with certainty, the network’s estimate of HD would better match the true HD, because evidence integration would be closer to Bayes-optimal. Here we show that this idea is correct.

Specifically, we measure the average accuracy of the network’s HD encoding (over many simulations), for both the Bayesian ring attractor and a conventional ring attractor. To model a conventional ring attractor, we use the same equations as we used for the Bayesian ring attractor, but we adjust the network connection strengths so that the bump amplitude decays to its stable baseline value very quickly (Figure 4a). Specifically, we strengthen both local recurrent excitatory connections (*w_sym_*) and global inhibition (*g*(*r_t_*)) while maintaining their balance, because their overall strengths are what controls the speed (*β*) of the bump’s return to its baseline amplitude (*κ*\*) in the regime near *κ*\*, assuming no change in the cell-intrinsic leak time constant *τ* (see Methods). With stronger overall connections, the bump amplitude decays to its stable baseline value more quickly. We then adjust the strength of global inhibition without changing the local excitation strength to maximize the accuracy of the network’s HD encoding; note that this changes *κ*\* but not *β*. This yields a conventional ring attractor where the bump amplitude is almost always fixed at a stable value (*κ*\*), with *κ*\* optimized for maximal encoding accuracy. Even after this optimization of the conventional ring attractor, it does not rival the accuracy of the Bayesian ring attractor. The Bayesian attractor performs consistently better, regardless of the amount of information available to the network, i.e., the level of certainty in the new HD observations (Figure 4b).

**Figure 4.**
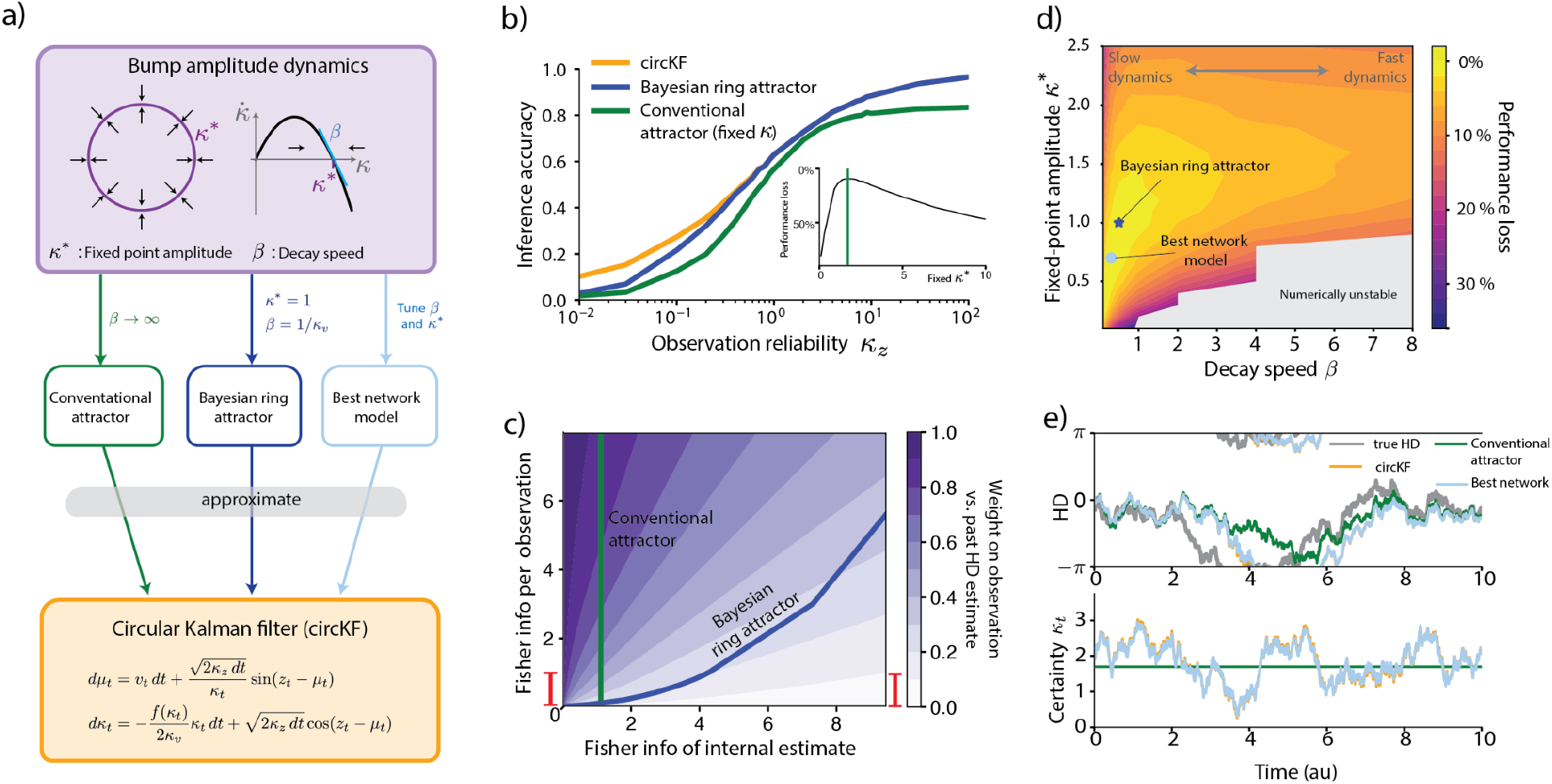
Ring attractors with slow dynamics approximate Bayesian inference. **a)** The ring attractor network in Eq. (3)) can be characterized by fixed point amplitude *κ*\* and decay speed *β*, which depend on the network connectivities. Thus, the network can operate in different regimes: a regime, where the bump amplitude is nearly constant (“Conventional attractor”), a regime where amplitude dynamics are tuned to implement a Bayesian ring attractor, or a regime with optimal performance (“Best network”, determined numerically). **b)** HD estimation performance as measured by inference accuracy (as defined by 1 – *circVar*, see Methods). For the “conventional” attractor, we chose *κ*\* to numerically maximize performance averaged across all levels of observation reliability, weighted by a prior *p*(*κ_z_*) on this reliability (see Methods). **c)** The weight with which a single observation contributes to the HD estimate varies with informativeness of both the HD observations and the current HD estimate. We here illustrate this for an HD observation that is orthogonal to the current HD estimate, resulting in the largest possible estimate change (|*z_t_* – *μ_t_*| = 90deg in Eq. (1)). The weight itself quantifies how much the observation impacts the HD estimate as a function of how informative this observation is (vertical axis, measured by Fisher information of a 10ms observation) and our certainty in the HD estimate (horizontal axis, also measured by Fisher information) before this observation. A weight of one implies that the observation replaces the previous HD estimate, whereas a weight of zero implies that the observation does not impact this estimate. The update weight of the Bayesian attractor is close to optimal (visually indistinguishable from the circKF; not shown here, but see Fig. S3), and forms a nonlinear curve through this parameter space. Fisher information per observation is directly related to the observation reliability *κ_z_*, and the vertical red bar shows the equivalent range of observation reliabilities, *κ_z_* ∈ [10^−2^,10^2^], shown in panel b. **d)** Overall inference performance loss (compared to a particle filter; performance measured by avg. inference accuracy, as in **b**, 0%: same average inference accuracy as particle filter, 100%: average. inference accuracy = 0), averaged across all levels of observation reliability (see Methods) as a function of the bump amplitude parameters *κ* * and *β*. The plot only shows performance loss for above-zero *β*’s and *κ*’s, as *β* → 0 or *κ* → 0 would cause the network’s required connectivity strengths to approach infinity. **e)** Simulated example trajectories of HD estimate/bump positions of HD estimate/bump positions (top) and certainties/bump amplitudes (bottom). To avoid cluttering, we are not showing the Bayesian ring attractor (visually indistinguishable from circKF and best network).

This performance difference arises because the conventional ring attractor does not keep track of the certainty associated with the current HD estimate. Ideally, the weight assigned to each HD observation depends on the current certainty of the current HD estimate, as well as the reliability of the observation itself (Figure 4c). A conventional ring attractor will assign more reliable observations a higher weight, but does not take into account the certainty of the current HD estimate. By contrast, the Bayesian ring attractor takes all these factors into account (Figure 5c).

To obtain more insight into the effect of bump decay speed (*β*) on network performance, we can also simulate many versions of our network with different values of *β*, which we generate by varying the overall strength of balanced local recurrent excitatory (*w_symmetric_*) and global inhibitory connections (*g*(*r_t_*)). We in turn vary the overall strength of global inhibition in order to find the best baseline bump amplitude (*κ*\*) for each value of *β*. The network with the best performance overall had a slow bump decay speed (low *β*), as expected (Figures 4d,e). As the bump decay speed *β* increased further, performance dropped. However, this could be partially mitigated by increasing baseline bump amplitude (*κ*\*) to prevent over-weighting of new observations.

We have seen that a slow bump decay (low *β*), i.e., the ability to deviate from the attractor state, is essential for uncertainty-related evidence weighting. That said, lower values of *β* are not always better. In the limit of very slow decay (*β* → 0), bump amplitude would grow so large that new HD observations have little influence, so that the network is nearly “blind” to visual landmarks. Conversely, in the limit of fast dynamics (*β* → ∞) the network is highly responsive to new observations; however, it also has almost no ability to weight those new observations relative to other observations in the recent past. In essence, *β* controls the speed of temporal discounting in evidence integration. Ideally, the bump decay speed *β* should be matched to the expected speed at which stored evidence becomes outdated and thus loses its value.

To summarize, we can frame the distinction between a conventional ring attractor and a Bayesian ring attractor as a difference in the speed of the bump’s decay to its stable state. In a conventional ring attractor, the bump decays quickly to its stable state, whereas in a Bayesian ring attractor, it decays slowly. Slow decay maximizes the accuracy of HD encoding because it allows the network to track its own internal certainty. Nonetheless, reasonable performance can be achieved even if the bump’s decay is fast, because a conventional ring attractor can still assign more informative observations a higher weight; it simply fails to assign the current HD estimate its proper weight.

### Tuning a biological ring attractor for Bayesian performance

Thus far, we have focused on model ring attractors with connection weights built from spatial cosine functions (Figure S3a), because this makes the mathematical treatment of these networks more tractable. However, this raises the question of whether a biological neural network can actually implement an approximation of the circular Kalman filter, even without these idealized connection weights. The most well-studied biological ring attractor network is the HD system of the fruit fly *Drosophila melanogaster* (Kim et al. 2017), and the detailed connections in this network have recently been mapped using large-scale electron microscopy connectomics (Hulse et al. 2021). We therefore asked whether the motifs from this connectomic data set -- and, by extension, motifs that could be found in any biological ring attractor network -- could potentially implement dynamic Bayesian inference.

To address this issue, we modeled the key cell types in this network (D. B. Turner-Evans et al. 2020; Scheffer et al. 2020; Hulse et al. 2021) (HD cells, angular velocity cells, and global inhibition cells), using connectome data to establish the patterns of connectivity between each cell type (SI text & Fig. S4b-f). We then analytically tuned the relative connection strengths between different cell types such that the dynamics of the bump parameters in the HD population implement an approximation of the circular Kalman filter. We also added a nonlinear element in the global inhibition layer, as this is required to approximate the circular Kalman filter. We found that this network achieves a HD encoding accuracy which is indistinguishable from that of our idealized Bayesian ring attractor network (Fig. S4g,h). Thus, even when we use connectome data to incorporate biological constraints on the network, the network is still able to implement dynamic Bayesian inference.

## Discussion

Uncertainty can affect navigation strategy (Merkle et al. 2006; Merkle and Wehner 2010), spatial cue integration (Cheng et al. 2007; Campbell et al. 2021), and spatial memory (Knight et al. 2014). This provides a motivation for understanding how uncertainty is represented in the neural networks that encode and store spatial variables for navigation. There is good reason to think that these networks are built around attractors. Thus, it is crucial to understand how attractors in general -- and ring attractors in particular -- might track uncertainty in spatial variables like head direction.

In this study, we have shown that a ring attractor can track uncertainty by operating in a dynamic regime away from its stable baseline states (its attractor states). In this regime, bump amplitude can vary, because we have tuned the local excitatory and global inhibitory connections in the ring attractor so they are relatively weak. By contrast, stronger overall connections produce a more conventional ring attractor that operates closer to its attractor states. Because the “Bayesian” ring attractor has a variable bump amplitude, bump amplitude grows when recent HD observations have been more reliable; in this situation, the network automatically ascribes more weight to its current estimate, relative to new evidence. Importantly, we have shown that nearly-optimal evidence weighting does not require exact tuning of the network connections. Indeed, even when we used connectome data to implement a network with realistic biological connectivity constraints, the network could still support near-optimal evidence weighting.

A key element of our approach is that bump amplitude is used to represent the internal certainty of the system’s HD estimate. In our framework, internal certainty determines the weight ascribed to new evidence, relative to past evidence. As such, the representation of internal certainty plays a crucial role in maximizing the accuracy of our Bayesian ring attractor. This stands in contrast to recent network models of the HD system that do not encode internal certainty, even though they assign more weight to more reliable HD observations, and less weight to less reliable HD observations (Sun et al. 2018; Sun et al. 2020). Notably, our network also automatically adjusts its cue integration weights to perform close-to-optimal Bayesian inference for HD observations of varying reliability. This differs from previous approaches (Esnaola-Acebes et al. 2021) that required hand-tuned weights.

Another important element of our approach was that we benchmarked our network model against a rigorous normative standard, the circular Kalman filter, which were derived analytically in (Kutschireiter et al. 2022) and here describe in terms of intuitive vector operations for the first time. Being able to rely on the circular Kalman filter was important because it allowed us to analytically derive the proper parameter values of our network model, so that the network’s estimate matched the estimate of an ideal observer. A remarkable property of the circular Kalman filter is that new HD observations will actually *decrease* certainty if they conflict strongly with the current estimate. This is not a property of a standard (non-circular) Kalman filter or a neural network designed to emulate it (Wilson and Finkel 2009). The power of conflicting evidence to decrease certainty is particular to the circular domain. Our Bayesian ring attractor network automatically reproduces this important aspect of the circular Kalman filter. Of course, the circular Kalman filter has applications beyond neural network benchmarking, as the accurate estimation of orientation or any other periodic variable has broad applications in the field of engineering.

In the brain’s HD system, the internal estimate of HD is based on not only HD observations (visual landmarks, etc.), but also angular velocity observations. The process of integrating these angular velocity observations over time is called angular path integration. Angular path integration is inherently noisy, and so uncertainty will grow progressively when HD observations are lacking. Our Bayesian ring attractor network is notable in explicitly treating angular path integration as a problem of probabilistic inference. Each angular velocity observation has limited reliability, and this causes the bump amplitude to decay in our network as long as HD observations are absent, in a manner that well-approximates the certainty decay of an ideal observer. In this respect, our network differs from previous investigations of ring attractors having variable bump amplitude (Carroll et al. 2014).

Our work makes several testable predictions. First, we predict that the HD system should contain the connectivity motifs required for a Bayesian ring attractor. Our analysis of *Drosophila* brain connectome data supports this idea; we expect similar network motifs to be present in the HD networks of other animals, such as that of mice (Peyrache et al. 2015; Ajabi et al. 2021), monkeys (Robertson et al. 1999), humans (Baumann and Mattingley 2010), or bats (Finkelstein et al. 2015). In the future, it will be interesting to determine whether synaptic inhibition in these networks is nonlinear, as predicted by our models.

Second, we predict that bump amplitude in the HD system should vary dynamically, with higher amplitudes in the presence of reliable external HD cues, such as salient visual landmarks. In particular, when bump amplitude is high, the bump’s position should be less sensitive to the appearance of new external HD cues. Notably, an experimental study from the mouse HD system provides some initial support for these predictions (Ajabi et al. 2021). This study found that the amplitude of population activity in HD neurons (what we call bump amplitude) increases in the presence of a reliable visual HD cue. Bump amplitude also varied spontaneously when all visual cues were absent (in darkness); intriguingly, when the bump amplitude was higher in darkness, the bump position was slower to change in response to the appearance of a visual cue, suggesting a lower sensitivity to the cue. In the future, more experiments will be needed to clarify the relationship between bump amplitude, certainty, and cue integration. In particular, it is puzzling that multiple studies (Zugaro et al. 2001; Shinder and Taube 2014; Seelig and Jayaraman 2015; Green et al. 2017; D. Turner-Evans et al. 2017; Ajabi et al. 2021) have found that bump amplitude increases with angular velocity, as higher angular velocities should not increase certainty.

In the future, more investigation will be needed to understand evidence accumulation on longer timescales. The circular Kalman filter is a recursive estimator: at each time step, it only considers the observer’s internal estimate from the previous time step, as well as the current observation of new evidence. However, when the environment changes, it would be useful to use a longer history of past observations (and past internal estimates) to readjust the weight assigned to the changing sources of evidence. Available data suggests that Hebbian plasticity can progressively strengthen the influence of the external sensory cues that are most reliably correlated with HD (Knierim et al.1998; Knight et al. 2014; Fisher et al. 2019; Kim et al. 2019). The interaction of Hebbian plasticity with attractor dynamics could provide a mechanism for extending statistical inference to longer timescales (Skaggs et al. 1994; Keinath et al. 2018; Milford et al. 2004; Mulas et al. 2016; Ocko et al. 2018; Page et al. 2014; Page and Jeffery 2018; Cope et al. 2017).

In summary, our work shows how ring attractors could implement dynamic Bayesian inference in the HD system. Our results have significance beyond the encoding of head direction -- e.g., they are potentially relevant for the grid cell ensemble, which appears to be organized around ring attractors even though it encodes linear rather than circular variables. Moreover, our models could apply equally to any brain system that needs to compute an internal estimate of a circular or periodic variable, such as visual object orientation (Li et al. 2021; van Bergen et al. 2015) or circadian time. More generally, our results demonstrate how canonical network motifs, like those common in ring attractor networks, can work together to perform close-to-optimal Bayesian inference, a problem with fundamental significance for neural computation.

## Acknowledgements

We thank Rachel Wilson for fruitful discussions and input throughout the whole research phase, and for her comments on the manuscript. We would like to thank Habiba Noamany for assisting us in navigating the neuprint database, and for informed comments on the manuscript. We would further like to thank Johannes Bill & Albert Chen for discussions and feedback on the manuscript, Philipp Reinhard for going on a typo hunt in the SI, and the entire Drugowitsch lab for valuable and insightful discussions.

The work was funded by the NIH (R34NS123819; J.D.), the James S. McDonnell Foundation (Scholar Award #220020462; J.D.), the Swiss National Science Foundation (grant numbers P2ZHP2 184213 and P400PB 199242; A.K.), and a Grant in the Basic and Social Sciences by the Harvard Medical School Dean’s Initiative award program (J.D.).

## Author contributions

Conceptualization, A.K., M.A.B., J.D.; Methodology, A.K., J.D.; Software, A.K.; Formal analysis, A.K., J.D.; Investigation, A.K, M.A.B., J.D.; Resources, J.D; Writing - Original Draft: A.K., J.D.; Writing - Review & Editing: A.K., M.A.B., J.D.; Visualization, A.K.; Supervision, J.D.; Funding Acquisition, A.K., J.D.

## Methods

### Ideal observer model: the circular Kalman filter

Our ideal observer model - the circular Kalman filter (circKF) [Kutschireiter et al., 2022] - performs dynamic Bayesian inference for circular variables. It computes the posterior probability of an unobserved (true) HD *ϕ_t_* ∈ [-*π*, *π*] at each point in time *t*, conditioned on a continuous stream of noisy angular velocity observations *v*_0:*t*_ = {*v*_0_, *v_dt_*,…*v_t_*} with 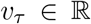, and HD observations *z*_0:*t*_ = {*z*_0_, *z_dt_*,…*z_t_*} with *z_τ_* ∈ [−*π*, *π*]. Specifically, we assume that these observations are generated from the true angular velocity 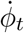 and HD *ϕ_t_*, corrupted by zero-mean noise at each point in time, via

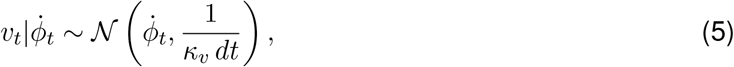

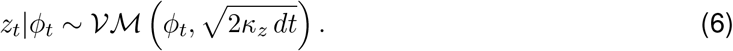

Here, 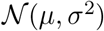 denotes a Gaussian with mean *μ* and variance *σ*^2^, 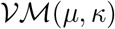 denotes a von Mises distribution of a circular random variable with mean *μ* and precision *κ*, and *κ_ν_* and *κ_z_* denote the precision of the angular velocity and HD observations, respectively. Note that as *dt* → 0, the precision of both angular velocity and HD observations approach 0, in line with the intuition that reducing a time step size *dt* results in more observations per unit time, which should be accounted for by less precision per observation to avoid “oversampling”. More formally, the square-root scaling of the HD observation precision with 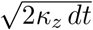 ensures that the Fisher information of the observations about the true HD scales linearly in time and *κ*_z_ in the continuum limit *dt* → 0 [Kutschireiter et al. 2022, Theorem 2]. The same applies to the dt^-1^ scaling of the Gaussian variance of the angular velocity observations, again achieving a Fisher information that scales linearly in time.

To support integrating information over time, the model assumes that current HD *ϕ_t_* depends on past HD *ϕ_t–dt_*. Specifically, in absence of further evidence, the model assumes that HD diffuses on a circle,

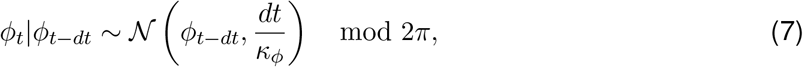

with a diffusion coefficient that decreases with *κ_ϕ_*. In Results, we assume *κ_ϕ_* → 0, implying that HD can change arbitrarily across consecutive time steps, which was sufficient to convey intuition into the algorithm’s workings. However, when simulating stochastic HD trajectories, we assume they evolve according to Eq. (7) with *κ_ϕ_* > 0, which needs to be accounted for when performing inference. Thus, we here assume a non-zero *K_ϕ_* for completeness and reproducibility.

The circKF in Eqs. (1) and (2) assumes that the posterior distribution over HD can be approximated by a von Mises distribution with time-dependent mean *μ_t_* and certainty *κ_t_*, i.e. 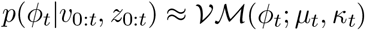. Such an approximation is justified if the posterior is sufficiently unimodal, and can, for instance, be compared to a similar approximation employed by extended Kalman filters for non-circular variables.

An alternative parametrization of the von Mises distribution to its mean *μ_t_* and precision *κ_t_*, are its natural parameters, *θ_t_* = (*κ_t_* cos *μ_t_*, *κ_t_* sin *μ_t_*)*^T^*. Geometrically, the natural parameters can be interpreted as the Cartesian coordinates of a “probability vector”, and (*μ_t_*, *κ_t_*) as its polar coordinates (Fig. 1b). As we show in the SI, the natural parameter parametrization makes including HD observations (Eq. (6)) in the circKF straightforward. In fact, it becomes a vector addition. In contrast, including angular velocity observations (Eq. (5)) is mathematically intractable, such that the circKF relies on an approximation method called projection filtering [Brigo et al., 1999] to find closed-form dynamic expressions for posterior mean and certainty (see [Kutschireiter et al., 2022] for technical details, and the SI for a more accessible description of the circKF).

Taken together, the circKF for the model specified by Eqs. (5)–(7) reads:

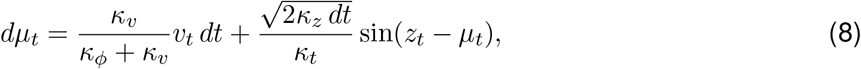

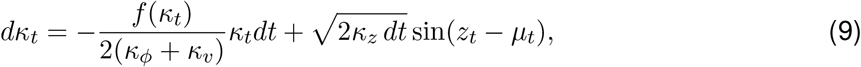

where *f*(*κ_t_*) is a monotonically increasing nonlinear function,

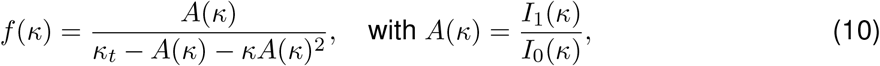

and *I*_0_(·) and *I*_1_(·) denote the modified Bessel functions of the first kind of order 0 and 1, respectively. Setting *κ_ϕ_* → 0 yields Eqs. (1) and (2). Importantly, a non-zero *κ_ϕ_* does not conceptually change the general vector operations we present in Fig. 1.

For a sufficiently large *κ* (i.e., high certainty), the nonlinearity *f*(*κ*) approaches the linear function, *f*(*κ*) → 2*κ* – 2. In our **quadratic approximation**, we thus replace the non-linearity by a quadratic decay:

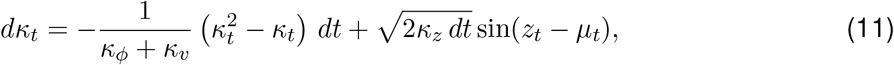

which well-approximates the circKF in the high certainty regime.

### Network model

We derived a rate-based network model that implements (approximations of) the circKF, by encoding the von Mises posterior parameters in activity 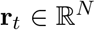 of a neural population with *N* neurons. Thereby, we focused on the simplest kind of network model that supports such an approximation, which is of the form:

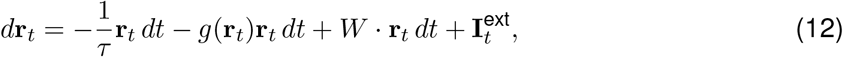

where *τ* is the cell-intrinsic leak time constant, 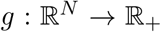 is a scalar nonlinearity, and the elements of r_t_ are assumed to be ordered by the respective neuron’s preferred HD, *ϕ*_1_,…, *ϕ_N_* (see Eq. (3)). We decomposed the recurrent connectivity matrix into 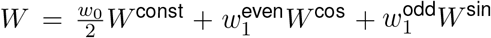, where *W*^const^ denotes a matrix with constant entries, and *W*^cos^ and *W*^sin^ refer to cosine- and sine-shaped connectivity profiles (Fig. 3d). In the main text we denote 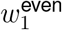 by *w*_sym_ and 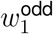 by *w*_asym_ to intuitively describe the connectivity pattern shown in Fig. 3d rather than the type of function (*even* and *odd*) or the Fourier component (subscript) that describes network connectivity in the continuous infinite-N limit. Irrespective of notation, the network’s circular symmetry makes the entries of these matrices only depend on the relative distance in preferred HD, and are given by 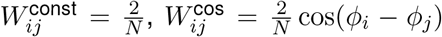, and 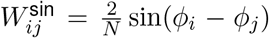. The scaling factor 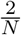 was chosen to facilitate matching our analytical results from the continuum network to the network structure outlined here. We further considered a cosine-shaped external input of the form 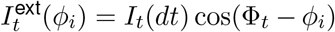 that is peaked around an input location Φ_*t*_. Here, *I_t_*(*dt*) denotes the input pattern in the infinitesimal time bin *dt*.

As described in Results, we assume the population activity r_*t*_ to encode the HD belief parameters *μ_t_* and *κ_t_* in the phase and amplitude of the activity’s first Fourier component. As we show in the SI, the described network dynamics thus lead to the following dynamics of the cosine-profile parameters *μ_t_* and *κ_t_*:

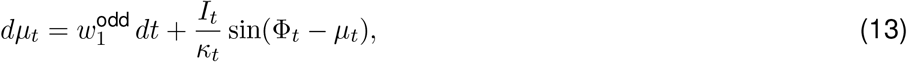

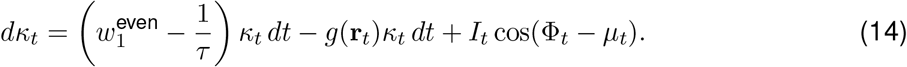

To derive these dynamics, we make the following three assumptions. First, we assume the network to be *rate-based*. Second, our analysis assumes a continuum of neurons, i.e. *N* → ∞. For numerical simulations, and the network description below, we used a finite-sized network of size N that corresponds to a discretization of the continuous network. Fig. S2 demonstrates only a very weak dependence of our results on the exact number of neurons in the network. Third, our analysis and simulations focused on the first Fourier mode of the bump profile, and is thus independent of the exact shape of the profile (as long as Eq. (3) holds).

### Network parameters for Bayesian inference

Having identified how the dynamics of the *μ_t_* and *κ_t_* encoded by the network (Eqs. (13) & (14)) depend on the network parameters, we now tuned these parameters to match these dynamics to those of the mean and certainty of the circKF (Eqs. (8) & (9)). Here, we first do so to achieve an exact match to the circKF, without the quadratic approximation. After that we describe the quadratic approximation that is used in the main text and leads to the Bayesian ring attractor network. Specifically, an exact match to the circKF requires the following network parameters:

- Odd recurrent connectivities are modulated by angular velocity observations, 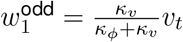, which shifts the activity profile without changing its amplitude [Skaggs et al., 1994, Zhang, 1996].
- HD observations *z_t_* are represented as the peak position Φ_*t*_ of a cosine-shaped external input whose amplitude is modulated by the reliability of the observation, i.e., 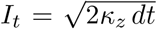. The inputs might contain additional Fourier modes (e.g., a constant baseline), but those do not affect the dynamics in Eqs. (13) and (14).
- The even component of the recurrent excitatory input needs to exactly balance the internal activity decay, i.e., 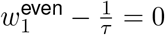.
- The decay nonlinearity is modulated by the reliability of the angular velocity observations, and is given by 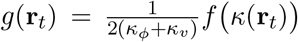, where *f*(·) equals the nonlinearity that governs the certainty decay in the circKF (Eq. (10)). This can be achieved, e.g., through interaction with an inhibitory neuron (or a pool of inhibitory neurons) with activation function *f*(·) that computes the activity bump’s amplitude *f*(*κ*(r_*t*_)).

A network with these parameters is *not* an attractor network, as its activity decays to zero in the absence of external inputs.

To arrive at the Bayesian ring attractor, we approximate the decay non-linearity by a quadratic approximation that takes the form 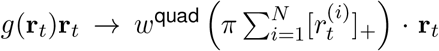, where [·]_+_ denotes the rectification nonlinearity. The resulting recurrent inhibition can be shown to be quadratic in the amplitude *κ_t_*, and has the further benefit of introducing an attractor state at a positive bump aplitude (see below). In the large population limit, *N* → ∞, this leads to the amplitude dynamics (see SI for derivation)

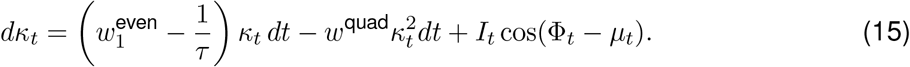

The dynamics of the phase μt does not depend on the form of *g*(·) and thus remains to be given by Eq. (13). If we set the network parameters to 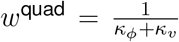 and 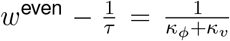, while sensory input, i.e. angular velocity *ν_t_* and HD observations *z_t_*, enter in the same way as before, the network implements the quadratic approximation to the circKF (Eqs. (8) & (11)).

### General ring-attractor networks with fixed point *κ*\* and decay speed *β*

In absence of HD observations (*I_t_* = 0), the amplitude dynamics in Eq. (15) has a stable **fixed point** at 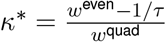 and no preferred phase, making it a ring-attractor network. Linearizing the *κ_t_* dynamics around this fixed points reveals that it is approached with **decay speed** 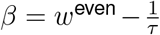. Therefore, we can tune the parameters to achieve a particular fixed point *κ*\* and decay speed *β* by setting *w*^even^ = *β* + 1/*τ* and 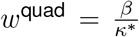. A large value of *β* requires increasing both *w*^even^ and *w*^quad^, and yields faster dynamics and thus indicates more rigid attractor dynamics. In the limit of *β* → ∞ the attractor becomes completely rigid in the sense that, upon any perturbation, it immediately moves back to its attractor state. In the main text we assume conventional ring attractors to operate close to this rigid regime. For the Bayesian ring attractor, we find *κ*\* = 1 and 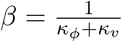. Further, in our simulations in Fig. 4, we explored network dynamics with a range of *κ*\* and *β* values by adjusting network parameters accordingly.

### Simulation details

#### Numerical integration

Our simulations in Figs. 3 and 4 used artificial data that matched the assumptions underlying our models. In particular, the ‘true’ HD *ϕ_t_* followed a diffusion on the circle, Eq. (7), and observations were drawn at each point in time from Eqs. (5) and (6). To simulate trajectories and observations, we used the Euler-Maruyama scheme [Kloeden and Platen, 2010], which supports the numerical integration of stochastic differential equations. Specifically, for a chosen discretization time step Δ*t*, this scheme is equivalent to drawing trajectories and observations from Eqs. (7), (5) and (6) directly while substituting dt → Δ*t*. The same time-discretization scheme was used to numerically integrate the SDEs of the circKF, Eqs (8) and (9), and its quadratic approximation, Eq. (11).

#### Performance measures

To measure performance, in Figs. 3f & 4b/d we computed the circular average distance [Mardia and Jupp, 2000] of the estimate *μ_T_* from the true HD **ϕ*_T_* at the end of a simulation of length *T* = 20 from *P =* 5^0^000 simulated trajectories by 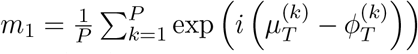. The absolute value of the imaginary-valued circular average, 0 ≤ |*m*_1_| ≤ 1 denotes an empirical accuracy (or ‘inference accuracy’), and thus measures how well the estimate *μ_T_* matches the true HD *ϕ_T_*. Here, a value of 1 denotes an exact match. The inference accuracy is related to the circular variance via Var_*circ*_ = 1 - |*m*_1_|. In Fig. S5, we provide histograms with samples *μ_T_* – *ϕ_T_* with different numerical values of |*m*_1_|, to provide some intuition for the spread of estimates for a given value of the performance measure.

We estimated performance through such averages for all HD observation reliabilities *κ_z_* in Figs. 3f & 4b. For the inset of Fig. 4b, and for Fig. 4d, we additionally performed a grid search over the fixed-point *κ*\* (inset of Fig. 4b), or both the fixed-point *κ*\* and of the decay speed *β* (Fig. 4d). For each setting of *κ*\* and *β* we assessed the performance by computing an average over this performance for a range of observation reliabilities *κ_z_*, weighted by how likely each observation reliability is a-priori assumed to be. The latter was specified by a log-normal prior, 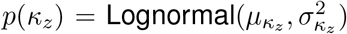), favouring intermediate reliabilitiy levels. We chose *μ_ν_z__* = 0.5 and 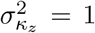 for the prior parameters, but our results did not strongly depend on this parameter choice. The performance loss shown in Fig. 4d also relied on such a weighted average across *κ_z_*’s for a particle filter benchmark (PF, see SI for details). The loss itself was then defined as 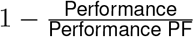.

#### Update weights for HD observations

In Fig. 4c, we computed the weight with which a single HD observation with |*z_t_* – *μ_t_*| = 90° changes the HD estimate. We defined this weight as the change in HD estimate, normalized by the value of the maximum possible change, 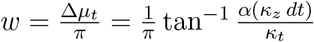. Here, *α*(*κ_z_ dt*) denotes a function that ensures a linear scaling of the Fisher information with sampling time step (see ref [Kutschireiter et al., 2022], Theorem 2, for details about this function). Thus, by design of the observation model, the Fisher information of a single observation with reliability *κ_z_* during a time interval Δ*t* is given by *I_zt_* (*ϕ_t_*) = *κ_z_* Δ*t*. We plot the weight as a function of the Fisher information of a single HD observation (how reliable is the observation?) and the Fisher information of the current HD estimate (how certain is the current estimate?), which is given by

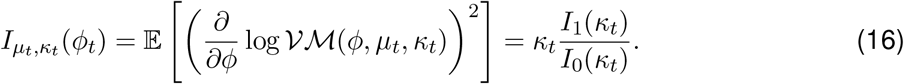

#### Details on numerical simulations

In our network simulations, we set the leak time constant *τ* to an arbitrary, but non-zero, value. Effectively, this resulted in a cosine-shaped activity profile. Note that by setting higher-order recurrent connectivities accordingly, other profile shapes could be realized, without affecting the validity of our analysis above. From the neural activity vector r_*t*_, we retrieved the natural parameters *θ_t_* with a decoder matrix *A* = (cos(*ϕ*), sin(*ϕ*))^*T*^, such that **θ**_*t*_ = *A* · r_*t*_, and subsequently computed the position of the bump by *ϕ_t_* = arctan2(*θ*_2_, *θ*_1_), and the encoded certainty (length of the population vector) by 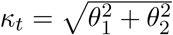.

In all our simulations, times are measured in units of inverse diffusion time constant *κ_ϕ_*, where we set *κ_ϕ_* = 1*s* for convenience. We used the following simulation parameters. For Fig. 3e, we used *κ_ν_* = 2, *κ_z_* = 10 (during ‘Visual cue’ period), *κ_z_* = 0 (during ‘Darkness’ period). For Figs. 3f & 4b/d we used *κ_ν_* = 1, *T* = 20, and averaged results over *P* = 5000 simulation runs. For Fig. 4e we used *κ_ν_* = 1, *κ_z_* = 1, *T* = 10. We used Δ*t* = 0.01 for all simulations.

Trajectory simulations and general analyses were performed on a MacBook Pro (Mid 2019) running 2.3 GHz 8-core Intel Core i9. Parameter scans were run on the Harvard Medical School O_2_ HPC cluster. For all our simulations, we used Python 3.9.1 with NumPy 1.19.2. Jupyter notebooks, Python scripts, and data to reproduce the figures will be made available upon acceptance of the manuscript.

## References

Ajabi, Zaki, Alexandra T. Keinath, Xue-Xin Wei, and Mark P. Brandon. 2021. “Population Dynamics of the Thalamic Head Direction System during Drift and Reorientation.” bioRxiv. https://doi.org/10.1101/2021.08.30.458266.

Amari, Shun-ichi. 1977. “Dynamics of Pattern Formation in Lateral-Inhibition Type Neural Fields.” Biological Cybernetics 27 (2): 77–87. https://doi.org/10.1007/BF00337259.

Baumann, O., and J. B. Mattingley. 2010. “Medial Parietal Cortex Encodes Perceived Heading Direction in Humans.” Journal of Neuroscience 30 (39): 12897–901. https://doi.org/10.1523/JNEUROSCI.3077-10.2010.

Beck, J. M., P. E. Latham, and A. Pouget. 2011. “Marginalization in Neural Circuits with Divisive Normalization.” Journal of Neuroscience 31 (43): 15310–19. https://doi.org/10.1523/JNEUROSCI.1706-11.2011.

Ben-Yishai, R., R. L. Bar-Or, and H. Sompolinsky. 1995. “Theory of Orientation Tuning in Visual Cortex.” Proceedings of the National Academy of Sciences 92 (9): 3844–48. https://doi.org/10.1073/pnas.92.9.3844.

Bergen, Ruben S van, Wei Ji Ma, Michael S Pratte, and Janneke F M Jehee. 2015. “Sensory Uncertainty Decoded from Visual Cortex Predicts Behavior.” Nature Neuroscience 18 (12): 1728–30. https://doi.org/10.1038/nn.4150.

Brigo, D., B. Hanzon, and F. Le Gland. 1999. “Approximate Nonlinear Filtering by Projection on Exponential Manifolds of Densities.” Bernoulli 5 (3): 495–534. https://doi.org/10.2307/3318714.

Campbell, Malcolm G., Alexander Attinger, Samuel A. Ocko, Surya Ganguli, and Lisa M. Giocomo. 2021. “Distance-Tuned Neurons Drive Specialized Path Integration Calculations in Medial Entorhinal Cortex.” Cell Reports 36 (10): 109669. https://doi.org/10.1016/j.celrep.2021.109669.

Carroll, Sam, Krešimir Josić, and Zachary P. Kilpatrick. 2014. “Encoding Certainty in Bump Attractors.” Journal of Computational Neuroscience 37 (1): 29–48. https://doi.org/10.1007/s10827-013-0486-0.

Cheng, Ken, Sara J. Shettleworth, Janellen Huttenlocher, and John J. Rieser. 2007. “Bayesian Integration of Spatial Information.” Psychological Bulletin.

Compte, A. 2000. “Synaptic Mechanisms and Network Dynamics Underlying Spatial Working Memory in a Cortical Network Model.” Cerebral Cortex 10 (9): 910–23. https://doi.org/10.1093/cercor/10.9.910.

Compte, Albert. 2006. “Computational and in Vitro Studies of Persistent Activity: Edging towards Cellular and Synaptic Mechanisms of Working Memory.” Neuroscience 139 (1): 135–51. https://doi.org/10.1016/j.neuroscience.2005.06.011.

Cope, Alex J., Chelsea Sabo, Eleni Vasilaki, Andrew B. Barron, and James A. R. Marshall. 2017. “A Computational Model of the Integration of Landmarks and Motion in the Insect Central Complex.” Edited by Paul Graham. PLOS ONE 12 (2): e0172325. https://doi.org/10.1371/journal.pone.0172325.

Dehaene, Guillaume P., Ruben Coen-Cagli, and Alexandre Pouget. 2021. “Investigating the Representation of Uncertainty in Neuronal Circuits.” PLOS Computational Biology 17 (2): e1008138. https://doi.org/10.1371/journal.pcbi.1008138.

Ermentrout, Bard. 1998. “Neural Networks as Spatio-Temporal Pattern-Forming Systems.” Reports on Progress in Physics 61 (4): 353–430. https://doi.org/10.1088/0034-4885/61/4/002.

Ernst, Marc O, and Martin S Banks. 2002. “Humans Integrate Visual and Haptic Information in a Statistically Optimal Fashion.” Nature 415 (6870): 429–33. https://doi.org/10.1038/415429a.

Esnaola-Acebes, Jose M., Alex Roxin, and Klaus Wimmer. 2021. “Bump Attractor Dynamics Underlying Stimulus Integration in Perceptual Estimation Tasks.” Preprint. Neuroscience. https://doi.org/10.1101/2021.03.15.434192.

Fetsch, Christopher R., Amanda H. Turner, Gregory C. DeAngelis, and Dora E. Angelaki. 2009. “Dynamic Reweighting of Visual and Vestibular Cues during Self-Motion Perception.” Journal of Neuroscience 29 (49): 15601–12. https://doi.org/10.1523/JNEUROSCI.2574-09.2009.

Finkelstein, Arseny, Dori Derdikman, Alon Rubin, Jakob N. Foerster, Liora Las, and Nachum Ulanovsky. 2015. “Three-Dimensional Head-Direction Coding in the Bat Brain.” Nature 517 (7533): 159–64. https://doi.org/10.1038/nature14031.

Fisher, Yvette E, Jenny Lu, Isabel D’Alessandro, and Rachel I Wilson. 2019. “Sensorimotor Experience Remaps Visual Input to a Heading-Direction Network.” Nature, no. December 2018 (November). https://doi.org/10.1038/s41586-019-1772-4.

Georgopoulos, Ap, Re Kettner, and Ab Schwartz. 1988. “Primate Motor Cortex and Free Arm Movements to Visual Targets in Three-Dimensional Space. II. Coding of the Direction of Movement by a Neuronal Population.” The Journal of Neuroscience 8 (8): 2928–37. https://doi.org/10.1523/JNEUROSCI.08-08-02928.1988.

Green, Jonathan, Atsuko Adachi, Kunal K. Shah, Jonathan D. Hirokawa, Pablo S. Magani, and Gaby Maimon. 2017. “A Neural Circuit Architecture for Angular Integration in Drosophila.” Nature 546 (7656): 101–6. https://doi.org/10.1038/nature22343.

Hansel, David, and Haim Sompolinsky. 1998. “Modeling Feature Selectivity in Local Cortical Circuits.” Methods in Neuronal Modeling: From Ions to Networks, 69.

Heinze, Stanley, Ajay Narendra, and Allen Cheung. 2018. “Principles of Insect Path Integration.” Current Biology: CB 28 (17): R1043–58. https://doi.org/10.1016/j.cub.2018.04.058.

Hopfield, J J. 1982. “Neural Networks and Physical Systems with Emergent Collective Computational Abilities.” Proceedings of the National Academy of Sciences 79 (8): 2554–58. https://doi.org/10.1073/pnas.79.8.2554.

Hulse, Brad K, Hannah Haberkern, Romain Franconville, Daniel B Turner-Evans, Shin-ya Takemura, Tanya Wolff, Marcella Noorman, et al. 2021. “A Connectome of the Drosophila Central Complex Reveals Network Motifs Suitable for Flexible Navigation and Context-Dependent Action Selection.” ELife 10 (October): e66039. https://doi.org/10.7554/eLife.66039.

Johnson, Adam, Kelsey Seeland, and A. David Redish. 2005. “Reconstruction of the Postsubiculum Head Direction Signal from Neural Ensembles.” Hippocampus 15 (1): 86–96. https://doi.org/10.1002/hipo.20033.

Kalman, R E. 1960. “A New Approach to Linear Filtering and Prediction Problems.” Transactions of the ASME Journal of Basic Engineering 82 (Series D): 35–45. https://doi.org/10.1115/1.3662552.

Kalman, R. E., and R. S. Bucy. 1961. “New Results in Linear Filtering and Prediction Theory.” Journal of Basic Engineering 83 (1): 95–108. https://doi.org/10.1115/1.3658902.

Keinath, Alexandra T, Russell A Epstein, and Vijay Balasubramanian. 2018. “Environmental Deformations Dynamically Shift the Grid Cell Spatial Metric.” Edited by Laura Colgin and Michael J Frank. ELife 7 (October): e38169. https://doi.org/10.7554/eLife.38169.

Kim, Sung Soo, Ann M Hermundstad, Sandro Romani, L F Abbott, and Vivek Jayaraman. 2019. “Generation of Stable Heading Representations in Diverse Visual Scenes.” Nature, no. December 2018: 1–6. https://doi.org/10.1038/s41586-019-1767-1.

Kim, Sung Soo, Hervé Rouault, Shaul Druckmann, and Vivek Jayaraman. 2017. “Ring Attractor Dynamics in the Drosophila Central Brain.” Science 356 (6340): 849–53. https://doi.org/10.1126/science.aal4835.

Kloeden, Peter E., and Eckhard Platen. 2010. Numerical Solution of Stochastic Differential Equations. Corr. 3. print. Applications of Mathematics 23. Berlin: Springer.

Knierim, James J., Hemant S. Kudrimoti, and Bruce L. McNaughton. 1998. “Interactions Between Idiothetic Cues and External Landmarks in the Control of Place Cells and Head Direction Cells.” Journal of Neurophysiology 80 (1): 425–46. https://doi.org/10.1152/jn.1998.80.1.425.

Knierim, James J., and Kechen Zhang. 2012. “Attractor Dynamics of Spatially Correlated Neural Activity in the Limbic System.” Annual Review of Neuroscience 35 (1): 267–85. https://doi.org/10.1146/annurev-neuro-062111-150351.

Knight, Rebecca, Caitlin E. Piette, Hector Page, Daniel Walters, Elizabeth Marozzi, Marko Nardini, Simon Stringer, and Kathryn J. Jeffery. 2014. “Weighted Cue Integration in the Rodent Head Direction System.” Philosophical Transactions of the Royal Society B: Biological Sciences 369 (1635): 20120512. https://doi.org/10.1098/rstb.2012.0512.

Knill, David C., and Alexandre Pouget. 2004. “The Bayesian Brain: The Role of Uncertainty in Neural Coding and Computation.” Trends in Neurosciences 27 (12): 712–19. https://doi.org/10.1016/j.tins.2004.10.007.

Kurz, Gerhard, Igor Gilitschenski, and Uwe D. Hanebeck. 2016. “Recursive Bayesian Filtering in Circular State Spaces.” IEEE Aerospace and Electronic Systems Magazine 31 (3): 70–87. https://doi.org/10.1109/MAES.2016.150083.

Kutschireiter, Anna, Luke Rast, and Jan Drugowitsch. 2022. “Projection Filtering with Observed State Increments with Applications in Continuous-Time Circular Filtering.” IEEE Transactions on Signal Processing, 1–1. https://doi.org/10.1109/TSP.2022.3143471.

Li, Hsin-Hung, Thomas C. Sprague, Aspen H. Yoo, Wei Ji Ma, and Clayton E. Curtis. 2021. “Joint Representation of Working Memory and Uncertainty in Human Cortex.” Neuron 109 (22): 3699–3712.e6. https://doi.org/10.1016/j.neuron.2021.08.022.

Lyu, Cheng, L. F. Abbott, and Gaby Maimon. 2022. “Building an Allocentric Travelling Direction Signal via Vector Computation.” Nature 601 (7891): 92–97. https://doi.org/10.1038/s41586-021-04067-0.

Ma, Wei Ji, Jeffrey M Beck, Peter E Latham, and Alexandre Pouget. 2006. “Bayesian Inference with Probabilistic Population Codes.” Nature Neuroscience 9 (11): 1432–38. https://doi.org/10.1038/nn1790.

Mardia, Kanti V., and Peter E. Jupp. 2000. Directional Statistics. John Wiley & Sons. https://doi.org/10.1002/9780470316979.

Merkle, Tobias, Markus Knaden, and Rüdiger Wehner. 2006. “Uncertainty about Nest Position Influences Systematic Search Strategies in Desert Ants.” Journal of Experimental Biology 209 (18): 3545–49. https://doi.org/10.1242/jeb.02395.

Merkle, Tobias, and Rüdiger Wehner. 2010. “Desert Ants Use Foraging Distance to Adapt the Nest Search to the Uncertainty of the Path Integrator.” Behavioral Ecology 21 (2): 349–55. https://doi.org/10.1093/beheco/arp197.

Milford, M.J., G.F. Wyeth, and D. Prasser. 2004. “RatSLAM: A Hippocampal Model for Simultaneous Localization and Mapping.” In IEEE International Conference on Robotics and Automation, 2004. Proceedings. ICRA ‘04. 2004, 1:403–408 Vol.1. https://doi.org/10.1109/ROBOT.2004.1307183.

Mulas, Marcello, Nicolai Waniek, and Jörg Conradt. 2016. “Hebbian Plasticity Realigns Grid Cell Activity with External Sensory Cues in Continuous Attractor Models.” Frontiers in Computational Neuroscience 10 (February). https://doi.org/10.3389/fncom.2016.00013.

Murray, Richard F., and Yaniv Morgenstern. 2010. “Cue Combination on the Circle and the Sphere.” Journal of Vision 10 (11): 15–15. https://doi.org/10.1167/10.11.15.

Ocko, Samuel A., Kiah Hardcastle, Lisa M. Giocomo, and Surya Ganguli. 2018. “Emergent Elasticity in the Neural Code for Space.” Proceedings of the National Academy of Sciences 115 (50). https://doi.org/10.1073/pnas.1805959115.

Page, Hector J. I., and Kate J. Jeffery. 2018. “Landmark-Based Updating of the Head Direction System by Retrosplenial Cortex: A Computational Model.” Frontiers in Cellular Neuroscience 12 (July): 191. https://doi.org/10.3389/fncel.2018.00191.

Page, Hector J. I., Daniel M. Walters, Rebecca Knight, Caitlin E. Piette, Kathryn J. Jeffery, and Simon M. Stringer. 2014. “A Theoretical Account of Cue Averaging in the Rodent Head Direction System.” Philosophical Transactions of the Royal Society B: Biological Sciences 369 (1635): 20130283. https://doi.org/10.1098/rstb.2013.0283.

Peyrache, Adrien, Marie M. Lacroix, Peter C. Petersen, and György Buzsáki. 2015. “Internally Organized Mechanisms of the Head Direction Sense.” Nature Neuroscience 18 (4): 569–75. https://doi.org/10.1038/nn.3968.

Piet, Alex T., Ahmed El Hady, Carlos D. Brody, Ahmed El Hady, and Carlos D. Brody. 2018. “Rats Adopt the Optimal Timescale for Evidence Integration in a Dynamic Environment.” Nature Communications 9 (1): 1–12. https://doi.org/10.1038/s41467-018-06561-y.

Rademaker, Rosanne L., Caroline H. Tredway, and Frank Tong. 2012. “Introspective Judgments Predict the Precision and Likelihood of Successful Maintenance of Visual Working Memory.” Journal of Vision 12 (13): 21. https://doi.org/10.1167/12.13.21.

Redish, A David, Adam N Elga, and David S Touretzky. 1996. “A Coupled Attractor Model of the Rodent Head Direction System.” Network: Computation in Neural Systems 7 (4): 671–85. https://doi.org/10.1088/0954-898X_7_4_004.

Robertson, Robert G., Edmund T. Rolls, Pierre Georges-François, and Stefano Panzeri. 1999. “Head Direction Cells in the Primate Pre-Subiculum.” Hippocampus 9 (3): 206–19. https://doi.org/10.1002/(SICI)1098-1063(1999)9:3<206::AID-HIPO2>3.0.CO;2-H.

Scheffer, Louis K, C Shan Xu, Michal Januszewski, Zhiyuan Lu, Shin-ya Takemura, Kenneth J Hayworth, Gary B Huang, et al. 2020. “A Connectome and Analysis of the Adult Drosophila Central Brain.” Edited by Eve Marder, Michael B Eisen, Jason Pipkin, and Chris Q Doe. ELife 9 (September): e57443. https://doi.org/10.7554/eLife.57443.

Seelig, Johannes D., and Vivek Jayaraman. 2015. “Neural Dynamics for Landmark Orientation and Angular Path Integration.” Nature 521 (7551): 186–91. https://doi.org/10.1038/nature14446.

Shinder, Michael E., and Jeffrey S. Taube. 2014. “Self-Motion Improves Head Direction Cell Tuning.” Journal of Neurophysiology 111 (12): 2479–92. https://doi.org/10.1152/jn.00512.2013.

Skaggs, William, James Knierim, Hemant Kudrimoti, and Bruce McNaughton. 1994. “A Model of the Neural Basis of the Rats Sense of Direction.” In Advances in Neural Information Processing Systems, edited by G. Tesauro, D. Touretzky, and T. Leen. Vol. 7. MIT Press. https://proceedings.neurips.cc/paper/1994/file/024d7f84fff11dd7e8d9c510137a2381-Paper.pdf.

Sun, Xuelong, Michael Mangan, and Shigang Yue. 2018. “An Analysis of a Ring Attractor Model for Cue Integration.” Lecture Notes in Computer Science (Including Subseries Lecture Notes in Artificial Intelligence and Lecture Notes in Bioinformatics) 10928 LNAI: 459–70. https://doi.org/10.1007/978-3-319-95972-6_49.

Sun, Xuelong, Shigang Yue, and Michael Mangan. 2020. “A Decentralised Neural Model Explaining Optimal Integration of Navigational Strategies in Insects.” Edited by Mani Ramaswami, Michael B Eisen, and Stanley Heinze. ELife 9 (June): e54026. https://doi.org/10.7554/eLife.54026.

Turner-Evans, Daniel B., Kristopher T. Jensen, Saba Ali, Tyler Paterson, Arlo Sheridan, Robert P. Ray, Tanya Wolff, et al. 2020. “The Neuroanatomical Ultrastructure and Function of a Biological Ring Attractor.” Neuron, September, S0896627320306139. https://doi.org/10.1016/j.neuron.2020.08.006.

Turner-Evans, Daniel, Stephanie Wegener, Hervé Rouault, Romain Franconville, Tanya Wolff, Johannes D Seelig, Shaul Druckmann, and Vivek Jayaraman. 2017. “Angular Velocity Integration in a Fly Heading Circuit.” ELife 6 (May): e23496. https://doi.org/10.7554/eLife.23496.

Wang, Xiao-Jing. 2001. “Synaptic Reverberation Underlying Mnemonic Persistent Activity.” Trends in Neurosciences 24 (8): 455–63. https://doi.org/10.1016/S0166-2236(00)01868-3.

Wilson, Robert, and Leif Finkel. 2009. “A Neural Implementation of the Kalman Filter.” Advances in Neural Information Processing Systems 22: 9.

Xie, Xiaohui, Richard H.R. Hahnloser, and H. Sebastian Seung. 2002. “Double-Ring Network Model of the Head-Direction System.” Physical Review E - Statistical Physics, Plasmas, Fluids, and Related Interdisciplinary Topics 66 (4): 9–9. https://doi.org/10.1103/PhysRevE.66.041902.

Zhang, Kechen. 1996. “Representation of Spatial Orientation by the Intrinsic Dynamics of the Head-Direction Cell Ensemble: A Theory.” The Journal of Neuroscience 16 (6): 2112–26. https://doi.org/10.1523/JNEUROSCI.16-06-02112.1996.

Zugaro, Michaël B., Eiichi Tabuchi, Céline Fouquier, Alain Berthoz, and Sidney I. Wiener. 2001. “Active Locomotion Increases Peak Firing Rates of Anterodorsal Thalamic Head Direction Cells.” Journal of Neurophysiology 86 (2): 692–702. https://doi.org/10.1152/jn.2001.86.2.692.

