## Supplementary material for "Bayesian inference in ring attractor networks": SI

PNAS
www.pnas.org

### Supplementary Information for

#### Bayesian inference in ring attractor networks

**Anna Kutschireiter, Melanie A. Basnak, and Jan Drugowitsch**

**Anna Kutschireiter,**
****

##### This PDF file includes:

Supplementary text
Figs. S1 to S5
References for SI reference citations

**5 Supplementary Figures** **19**

#### Supporting Information Text

#### 1. Circular Kalman filtering

Here, we present a derivation of the circular Kalman filter (circKF), which we use as an ideal observer model in the main text. The following derivation's main purpose is to provide the reader with some intuition behind the formalism, such that it uses a discrete-time approximation, followed by taking the continuous-time limit. For a mathematically-rigorous, continuous-time derivation of the circKF, please consult (1).

**A. Generative model.** Assuming time to be discretized in steps of $dt$, the overall goal is to derive an online estimator for the unobserved true head direction (HD) $\phi_t \in [-\pi, \pi]$ at each point in time $t$, conditioned on a continuous stream of noisy angular velocity observations $V_t = \{v_0, v_{dt}, \dots v_t\}$ (in the main text denoted $v_{0:t}$) with $v_\tau \in \mathbb{R}$ and HD observations $Z_t = \{z_0, z_{dt}, \dots z_t\}$ (in the main text denoted $z_{0:t}$) with $z_\tau \in [-\pi, \pi]$. We assume that these observations are generated from the (true) angular velocity $\dot{\phi}_t = \frac{\phi_t - \phi_{t-dt}}{dt}$ and HD $\phi_t$, respectively, and are corrupted by zero-mean noise at each point in time:

$$p(v_t|\phi_t, \phi_{t-dt}) = \mathcal{N}\left(v_t; \frac{\phi_t - \phi_{t-dt}}{dt}, \frac{1}{\kappa_v\, dt}\right), \tag{S1}$$

$$p(z_t|\phi_t) = \mathcal{VM}\left(z_t; \phi_t, \sqrt{2\kappa_z\, dt}\right), \tag{S2}$$

where $\mathcal{VM}(\varphi; \mu, \kappa) = \frac{e^{\kappa\cos(\varphi-\mu)}}{2\pi I_0(\kappa)}$ denotes the von Mises distribution of a circular random variable $\varphi$ with mean $\mu$ and precision $\kappa$. $\kappa_v$ and $\kappa_z$ refer to the precision of the angular velocity and HD observations, respectively. The noise scaling in the HD observations, $\sqrt{2\kappa_z\, dt}$, is chosen such that that the Fisher information about the HD of a single observation scales linearly with the sampling time step $dt$ in the limit $dt \to 0$ (1, Theorem 2).

We further assume that HD $\phi_t$ follows a diffusion on the circle, which serves as a dynamic prior over HD in terms of a transition density:

$$p(\phi_t|\phi_{t-dt}) \sim \mathcal{N}\left(\phi_t; \phi_{t-dt}, \frac{dt}{\kappa_\phi}\right) \mod 2\pi, \tag{S3}$$

Here, $\kappa_\phi \geq 0$ is related to the inverse diffusion constant: a large $\kappa_\phi$ implies limited diffusion and an almost-stationary stochastic process. In this case, past observations are generally highly informative about the current HD estimate. A small $\kappa_\phi$ implies that HD is most likely to change significantly from one time step to the next, indicating that past observations only provide limited information about our current HD.

**B. Discrete-time Bayesian filtering.** Given the posterior $p(\phi_{t-dt}|Y_{t-dt}, Z_{t-dt})$ at some previous time-step $t - dt$, we compute the posterior at the current time step $t$ using the conditional dependencies of the model and Bayes' theorem:

$$\begin{aligned} p(\phi_t|V_t, Z_t) &\propto_{\phi_t} p(z_t|\phi_t)p(\phi_t|V_t, Z_{t-dt}) \\ &= p(z_t|\phi_t)\int d\phi_{t-dt}\, p(\phi_t|\phi_{t-dt}, v_t)p(\phi_{t-dt}|Z_{t-dt}, V_{t-dt}). \end{aligned} \tag{S4}$$

This equation offers a way to *recursively* compute the current posterior density from the previous one, by taking two distinct steps: the so-called prediction and update step. The *prediction step* is a convolution between the previous posterior and the transition density $p(\phi_t|\phi_{t-dt}, v_t)$, as implemented by the above integral. It tells us how the posterior is expected to evolve in a single time step when only observing angular velocity information, but no HD observations, are present, resulting in the prediction density $p(\phi_t|V_t, Z_{t-dt})$. Note that the angular velocity observations $v_t$ enter this step through the effective transition probability $p(\phi_t|\phi_{t-dt}, v_t)$. In the *update step*, we multiply the result of the prediction step with the HD observation likelihood $p(z_t|\phi_t)$. Intuitively, this step can be understood as Bayesian cue integration between the prediction density and the HD observations.

In general, we will not be able to solve Eq. [S4] in closed form* for continuous variables like HD. We thus have to introduce approximations of $p(\phi_t|V_t, Z_t)$ that allow us to consistently perform prediction and update steps. Specifically, as one of the simplest choices for unimodal probability distributions for circular variables, we chose to approximate the posterior by a von Mises distribution,

$$p(\phi_t|V_t, Z_t) \approx \mathcal{VM}(\phi_t; \mu_t, \kappa_t). \tag{S5}$$

By using this approximation, the estimation task reduces to having to find evolution equations, conditioned on angular velocity observations $v_t$ and HD observations $z_t$, for the two parameters $\mu_t$ and $\kappa_t$, which are sufficient to fully specify the posterior distribution. In what follows, we will consider the effect of angular velocity observations and HD observations on the two parameters separately.

*In fact, a closed-form solution is almost never achievable for continuous state-spaces. One of the few cases where it is is when prediction and update steps are linear Gaussians, in which case Eq. [S4] yields the Kalman filter.

***B.1. Angular velocity observations.*** In Eq. [S4], angular velocity observations enter through a modified transition density $p(\phi_t|\phi_{t-dt}, v_t)$, which can be computed using Bayes' theorem:

$$p(\phi_t|v_t, \phi_{t-dt}) \propto_{\phi_t} p(v_t|\phi_t, \phi_{t-dt})p(\phi_t|\phi_{t-dt}). \tag{S6}$$

The modified transition probability is again a Gaussian, as can be seen from its logarithm being quadratic in $\phi_t$,

$$\begin{aligned} -\log p(\phi_t|v_t, \phi_{t-dt}) &= \frac{\kappa_v dt}{2}\left(v_t - \frac{\phi_t - \phi_{t-dt}}{dt}\right)^2 + \frac{\kappa_\phi}{2dt}\left(\phi_t - \phi_{t-dt}\right)^2 + \mathcal{R} \\ &= \frac{1}{2}\frac{\kappa_v + \kappa_\phi}{dt}\left(\phi_t - \phi_{t-dt}\right)^2 - \frac{\kappa_v}{dt}\left(\phi_t - \phi_{t-dt}\right) v_t dt + \mathcal{R} \\ &= \frac{1}{2}\frac{\kappa_v + \kappa_\phi}{dt}\left(\phi_t - \left(\phi_{t-dt} + \frac{\kappa_v}{\kappa_v + \kappa_\phi} v_t dt\right)\right)^2 + \mathcal{R}, \end{aligned} \tag{S7}$$

where terms independent of $\phi_t$, collectively denoted by $\mathcal{R}$, can be absorbed in the normalization. Hence, the modified transition probability reads:

$$p(\phi_t|v_t, \phi_{t-dt}) = \mathcal{N}\left(\phi_t; \phi_{t-dt} + \frac{\kappa_v}{\kappa_\phi + \kappa_v} v_t\, dt, \frac{dt}{\kappa_\phi + \kappa_v}\right) \quad \text{mod } 2\pi. \tag{S8}$$

Together with the assumption that the posterior of the last time step, $p(\phi_{t-dt}|V_{t-dt}, Z_{t-dt})$, is given by a von Mises distribution with mean $\mu_{t-dt}$ and precision $\kappa_{t-dt}$, we can write down the expression for the prediction density $p(\phi_t|V_t, Z_{t-1})$ (cf. first line in Eq. [S4]):

$$\begin{aligned} p(\phi_t|V_t, Z_{t-dt}) &= \int_{-\pi}^{\pi} d\phi_{t-dt}\, p(\phi_t|v_t, \phi_{t-dt})p(\phi_{t-dt}|Z_{t-dt}, V_{t-dt}) \\ &= \int_{-\pi}^{\pi} d\phi_{t-dt}\, \mathcal{N}\left(\phi_t; \phi_{t-dt} + \frac{\kappa_v}{\kappa_\phi + \kappa_v} v_t\, dt, \frac{dt}{\kappa_\phi + \kappa_v}\right) \mathcal{VM}\left(\phi_{t-dt}; \mu_{t-dt}, \kappa_{t-dt}\right). \end{aligned} \tag{S9}$$

Unfortunately, there is no closed-form solution for this integral. To approximate the prediction density $p(\phi_t|V_t, Z_{t-1})$ at each moment in time by a von Mises density $\mathcal{VM}(\phi_t; \tilde{\mu}_t, \tilde{\kappa}_t)$, we will use a more sophisticated approximation method, namely a projection filter (2). Such a filter ensures that this approximation is optimal by minimizing the infinitesimal Kullback-Leibler divergence at each moment in time. The technical details can be found in (1), and in this SI we limit ourselves to giving the final result:

$$d\mu_t = \frac{\kappa_v}{\kappa_v + \kappa_\phi} v_t\, dt, \tag{S10}$$

$$d\kappa_t = -\frac{f(\kappa_t)}{2(\kappa_v + \kappa_\phi)} \kappa_t\, dt. \tag{S11}$$

Here, the decay of the certainty $\kappa_t$ is governed by the nonlinear function

$$f(\kappa_t) = \frac{A(\kappa_t)}{\kappa_t - A(\kappa_t) - \kappa A(\kappa_t)^2}, \quad \text{with } A(\kappa_t) = \frac{I_1(\kappa_t)}{I_0(\kappa_t)}, \tag{S12}$$

where $I_0(\cdot)$ and $I_1(\cdot)$ denote the modified Bessel functions of the first kind of order 0 and 1. This function takes care of the fact that the true HD $\phi_t$ follows a diffusion on the circle, which becomes particularly relevant for small values of $\kappa_t$. In particular, $f(\kappa_t) \approx 1$ for small $\kappa_t$ and $f(\kappa_t) \approx 2\kappa_t - 2$ for large $\kappa_t$, indicating that the decay is asymptotically quadratic.

***B.2. HD observations.*** Angular-valued HD observations $z_t$ are integrated by multiplying the observation likelihood $p(z_t|\phi_t)$ with the prediction density $p(\phi_t|V_t, Z_{t-dt})$. If the prediction density is also von Mises (which is the assumption above), this cue integration is closed:

$$\begin{aligned} p(\phi_t|z_t, dy_t) &= \mathcal{VM}(z_t; \phi_t, \sqrt{2\kappa_z\, dt}) \cdot \mathcal{VM}(\phi_t; \tilde{\mu}_t, \tilde{\kappa}_t) \\ &\propto \exp\left(\begin{pmatrix}\cos\phi_t \\ \sin\phi_t\end{pmatrix}^\top \cdot \left(\sqrt{2\kappa_z\, dt}\begin{pmatrix}\cos z_t \\ \sin z_t\end{pmatrix} + \tilde{\kappa}_t \begin{pmatrix}\cos\tilde{\mu}_t \\ \sin\tilde{\mu}_t\end{pmatrix}\right)\right) \end{aligned} \tag{S13}$$

$$\stackrel{!}{=} \exp\left(\begin{pmatrix}\cos\phi_t \\ \sin\phi_t\end{pmatrix}^\top \cdot \kappa_t \begin{pmatrix}\cos\mu_t \\ \sin\mu_t\end{pmatrix}\right). \tag{S14}$$

Thus, the natural parameters of the posterior distribution, $\boldsymbol{\theta}_t = (\theta_1, \theta_2) = (\kappa_t \cos\mu_t, \kappa_t \sin\mu_t)^\top$, can be written as the sum of the natural parameters of the prediction density and the likelihood[†]:

$$\boldsymbol{\theta}_t = \tilde{\boldsymbol{\theta}}_t + \sqrt{2\kappa_z\, dt}\begin{pmatrix}\cos z_t\\ \sin z_t\end{pmatrix} \quad \text{[S15]}$$

$$d\boldsymbol{\theta}_t = \boldsymbol{\theta}_t - \tilde{\boldsymbol{\theta}}_t = \sqrt{2\kappa_z\, dt}\begin{pmatrix}\cos z_t\\ \sin z_t\end{pmatrix}. \quad \text{[S16]}$$

The updates of the parameters $\mu_t$ and $\kappa_t$ of the von Mises distribution due to the observation $z_t$ are obtained by transforming the update of $\boldsymbol{\theta}_t$ to polar coordinates:

$$d\mu_t^{\text{update}} = d\arctan 2\,(\theta_2, \theta_1) = \frac{\sqrt{2\kappa_z\, dt}}{\kappa_t}\sin(z_t - \mu_t) \quad \text{[S17]}$$

$$d\kappa_t^{\text{update}} = d\sqrt{\theta_1^2 + \theta_2^2} = \sqrt{2\kappa_z\, dt}\cos(z_t - \mu_t). \quad \text{[S18]}$$

***B.3. The circular Kalman filter.*** In the continuum limit $dt \to 0$, we do not distinguish between the parameters of the prediction density, $\tilde{\mu}_t$ and $\tilde{\kappa}_t$, and that of the posterior density, $\mu_t$ and $\kappa_t$. The circKF equations result from taking the prediction and update steps simultaneously, thereby combining Eq. [S10] with Eq. [S17] for the mean dynamics, and Eq. [S11] with Eq. [S18] for the precision dynamics:

$$d\mu_t = \frac{\kappa_v}{\kappa_\phi + \kappa_v} v_t\, dt + \frac{\sqrt{2\kappa_z\, dt}}{\kappa_t}\sin(z_t - \mu_t), \quad \text{[S19]}$$

$$d\kappa_t = -\frac{f(\kappa_t)}{2(\kappa_\phi + \kappa_v)}\kappa_t dt + \sqrt{2\kappa_z\, dt}\cos(z_t - \mu_t). \quad \text{[S20]}$$

Here, we adhered to expressing these equations in terms of their infinitesimal difference, $d\mu_t$ and $d\kappa_t$, instead of a differential equation. This is a standard way to express stochastic differential equations (SDEs), which makes it more straightforward to deal with the non-linear time scaling of the HD observations $z_t$.

***B.4. The quadratic approximation of the circular Kalman filter.*** If $\kappa_t$ is sufficiently large, the nonlinearity $f(\kappa_t)$ can be approximated by a linear function, $f(\kappa_t) \approx 2\kappa_t - 2$, such that the decay in Eq. [S20] becomes quadratic:

$$d\kappa_t \approx -\frac{1}{\kappa_\phi + \kappa_v}\left(\kappa_t^2 - \kappa_t\right) dt + \sqrt{2\kappa_z\, dt}\cos(z_t - \mu_t). \quad \text{[S21]}$$

We use this approximation when implementing the Bayesian ring attractor network.

**C. Coordinate transforms [Technical].** The von Mises distribution can be parametrized by its mean and precision parameters, $\mu$ and $\kappa$, or in terms of its natural parameters, $\boldsymbol{\theta} = (\theta_1, \theta_2)^\top = (\kappa\cos\mu, \kappa\sin\mu)^\top$. These two parametrizations are perfectly equivalent, and can be thought of as the polar and Cartesian coordinates of a vector, respectively. Except when $\kappa = 0$, which we assume to never occur, we can go back and forth between these representations by performing a coordinate transformation.

For the neural network we describe further below, it is easier to decode $\boldsymbol{\theta}$ than $\mu$ and $\kappa$ from neural population activity. Thus, it is useful to express the circular Kalman filter as SDEs for $\boldsymbol{\theta}$. Unfortunately, we cannot simply find these SDEs by applying a coordinate transform to Eqs. [S19] and [S20]. Technically speaking, since the angular velocity observations $v_t$ follow a stochastic process, we have to take into account second-order derivatives, which is called Itô's lemma in stochastic calculus (see (3) for an introduction). As we will here show in a slightly technical argument, using stochastic instead of ordinary calculus explains why we need an additional decay term in the network implementation in Sec. 3 that would not arise from a simple coordinate transform. Understanding this argument is not required for understanding our general theory and network implementation, and thus can safely be skipped.

First, we express the generative model in Eqs. [S3] and [S1] in terms of their equivalent Itô stochastic differential equations (SDEs). Defining the infinitesimal increment $du_t := v_t\, dt$, the SDEs read:

$$d\phi_t = \frac{1}{\sqrt{\kappa_\phi}} dW_t \quad \text{[S22]}$$

$$du_t = d\phi_t + \frac{1}{\sqrt{\kappa_v}} dV_t, = \frac{1}{\sqrt{\kappa_\phi}} dW_t + \frac{1}{\sqrt{\kappa_v}} dV_t, \quad \text{[S23]}$$

where $dW_t \in \mathbb{R} \sim \mathcal{N}(0, dt)$ and $dV_t \in \mathbb{R} \sim \mathcal{N}(0, dt)$ are uncorrelated scalar-valued Brownian motion processes with $dW_t\, dV_t = 0$. Since the variance of Brownian motion processes grows linearly in time, we have that $(dW_t)^2 = dt$, $(dV_t)^2 = dt$, and thus $(du_t)^2 = \left(\frac{1}{\kappa_\phi} + \frac{1}{\kappa_v}\right) dt$. The second equality in Eq. [S23] tells us that whenever angular velocity observations are drawn from the 'true' generative model in Eq. [S1], they automatically inherit the noise of the process that was used to generate $\phi_t$.

[†] This is not too surprising, as it is well known that in exponential family distributions these update steps boil down to adding up the natural parameters.

Itô's lemma tells us how to perform a variable transformation from a stochastic process $x_t$, which is governed by an Itô SDE, to another stochastic process $y_t = g(x_t)$:

$$dy_t = dg(x_t) = \left.\frac{\partial g(x)}{\partial x}\right|_{x=x_t} dx_t + \frac{1}{2}\left.\frac{\partial^2 g(x)}{\partial x^2}\right|_{x=x_t} (dx_t)^2. \tag{S24}$$

Thus, we can use Itô's lemma to transform the dynamics of $\mu_t$ and $\kappa_t$ in Eqs. [S10] and [S11] to the dynamics of the natural parameters of the von Mises distribution. Note that, since the dynamics of $\kappa_t$ are independent of the angular velocity observations, Eq. [S11] is deterministic with $(d\kappa_t)^2 = 0$:

$$\begin{aligned} d\boldsymbol{\theta}_t = d\left[\kappa_t \begin{pmatrix}\cos\mu_t\\ \sin\mu_t\end{pmatrix}\right] &= \begin{pmatrix}\cos\mu_t\\ \sin\mu_t\end{pmatrix} d\kappa_t + \kappa_t \begin{pmatrix}-\sin\mu_t\\ \cos\mu_t\end{pmatrix} d\mu_t + \frac{1}{2}\kappa_t \begin{pmatrix}-\cos\mu_t\\ -\sin\mu_t\end{pmatrix} (d\mu_t)^2 \\ &= -\frac{f(\kappa_t)}{2(\kappa_\phi+\kappa_v)}\kappa_t \begin{pmatrix}\cos\mu_t\\ \sin\mu_t\end{pmatrix} dt + \frac{\kappa_t\kappa_v}{\kappa_\phi+\kappa_v}\begin{pmatrix}-\sin\mu_t\\ \cos\mu_t\end{pmatrix} du_t - \frac{1}{2}\begin{pmatrix}\theta_1\\ \theta_2\end{pmatrix}\frac{\kappa_v^2}{(\kappa_v+\kappa_\phi)^2}(du_t)^2 \\ &= -\frac{1}{2}\frac{f(\kappa_t)}{\kappa_v+\kappa_\phi}\boldsymbol{\theta}_t\, dt - \frac{1}{2}\frac{\kappa_v/\kappa_\phi}{\kappa_v+\kappa_\phi}\boldsymbol{\theta}_t\, dt + \frac{\kappa_v}{\kappa_v+\kappa_\phi}\begin{pmatrix}0 & -1\\ 1 & 0\end{pmatrix}\boldsymbol{\theta}_t\, du_t. \end{aligned} \tag{S25}$$

Here, the additional decay term $-\frac{1}{2}\frac{\kappa_v/\kappa_\phi}{\kappa_v+\kappa_\phi}\boldsymbol{\theta}_t dt$ arises from the stochastic nature of the increment process $u_t$.

Since HD observations $z_t$ are added on the level of natural parameters (cf. Eq. [S16]), these can be included in a straightforward manner, yielding the circular Kalman filter in its natural parameter form:

$$d\boldsymbol{\theta}_t = -\frac{1}{2}\frac{f(\kappa_t)+\kappa_v/\kappa_\phi}{\kappa_v+\kappa_\phi}\boldsymbol{\theta}_t\, dt + \frac{\kappa_v}{\kappa_v+\kappa_\phi}\begin{pmatrix}0 & -1\\ 1 & 0\end{pmatrix}\boldsymbol{\theta}_t\, du_t + \sqrt{2\kappa_z\, dt}\begin{pmatrix}\cos z_t\\ \sin z_t\end{pmatrix}. \tag{S26}$$

**D. Numerical benchmarks.** As described above, the circKF approximates the posterior at each point in time by a von Mises distribution, and thus is itself an approximate algorithm. To compare its performance, and that of the Bayesian ring attractor to the truly best filtering performance for the assumed generative model, we additionally used a Bootstrap particle filter, which is exact in the limit of an infinite number of particles. Here, we first outline the algorithm itself, and then discuss how we assess filtering performance in general, to compare performance across algorithms.

***D.1. Bootstrap particle filter.*** As a numerical benchmark, we used a Sequential Importance Sampling/Resampling particle filter (4) (SIS-PF; member of the family of Bootstrap particle filters) that we modified to be applicable to angular velocity observations. Here, we briefly outline the numerical implementation of the SIS-PF for our particular filtering problem, and refer the reader to more specialized literature for derivation and convergence results (e.g., in (4, 5)).

The principle behind particle filters is that they provide a weighted empirical estimate of the posterior distribution,

$$p\left(\phi_t | V_t, Z_t\right) \approx \sum_{i=1}^{N} w_t^{(i)} \delta(\phi_t - \varphi_t^{(i)}), \tag{S27}$$

where we refer to $w_t^{(i)}$ as the importance weight of the $i$-th particle with position $\varphi_t^{(i)}$. Weighted particle filters are asymptotically exact, i.e. they provide us with the best possible inference performance in the limit of infinitely many particles $N \to \infty$. At each discrete time step, the $N$ particles in the SIS-PF are propagated according to the proposal density $\pi$, which we chose to correspond to the modified transition density in Eq. [S8]:

$$\begin{aligned} &\pi\left(\varphi_t^{(j)} | \varphi_{t-\Delta t}^{(j)}, v_t\right) \\ &\quad = \mathcal{N}\left(\varphi_t^{(j)}; \varphi_{t-\Delta t}^{(j)} + \frac{\kappa_v}{\kappa_v+\kappa_\varphi} v_t\, \Delta t, \frac{\Delta t}{\kappa_\varphi+\kappa_v}\right) \mod 2\pi. \end{aligned} \tag{S28}$$

Subsequently, each particle $j$ is weighted at each time step according to how well the proposed particle distribution fits to the HD observation $z_t$. This is equivalent to multiplying the previous weight with the observation likelihood (Eq. [S2]):

$$w_t^{(i)} = w_{t-\Delta t}^{(i)} \cdot \mathcal{VM}\left(z_t; \varphi_t^{(i)}, \sqrt{2\kappa_z \Delta t}\right). \tag{S29}$$

Lastly, the particles are re-weighted such that the importance weights sum to 1, $\sum_i w_t^{(i)} = 1$:

$$w_t^{(i)} \leftarrow \frac{w_t^{(i)}}{\sum_j w_t^{(j)}} \tag{S30}$$

In our simulations, we used $N = 10^3$ particles, which is sufficient if HD observations are present.

Mean $\mu_t$ and precision $r_t \in [0, 1]$ of the filtering distribution approximated by the SIS-PF can be determined at each time step according to a weighted average on the circle, i.e. the first circular moment:

$$r_t \exp(i\mu_t) = \sum_{j=1}^{N} w_t^{(j)} \exp\left(i\varphi_t^{(j)}\right). \qquad \text{[S31]}$$

***D.2. HD tracking performance measures.*** In the main text, we quantified HD tracking performance by estimating the absolute value of the circular average distance between the estimate $\mu_T$ at the end of the trial (using the mean of the filter posterior, which is the filter's best guess), and the true HD $\phi_T$, averaged across $P$ simulations with different noisy observation sequences, $v_0, \dots, v_T$ and $z_0, \dots, z_T$:

$$m_1 = \frac{1}{P} \sum_{k=1}^{P} \exp\left(i\left(\mu_T^{(k)} - \phi_T^{(k)}\right)\right). \qquad \text{[S32]}$$

Here, $m_1$ is a complex number, and HD tracking performance corresponds to its absolute value, $|m_1|$ (larger = better / more accurate). Note that this absolute value is one minus the circular variance of the error. As this variance is bounded by zero and one, zero variance implies a performance of $|m_1| = 1$, and maximum variance of one implies a performance of $|m_1| = 0$. To get a sense of how estimates $\mu_T$ are distributed around the true HD $\phi_T$ for a given value of $|m_1|$, we provide representative histograms in Fig. S5.

#### 2. Neural encoding example: encoding of the von Mises distribution with a linear probabilistic population code

In the main text, we assume a bump-like encoding of the HD estimate whose bump amplitude is scaled by the encoded certainty $\kappa_t$. This implies that the amplitude of the first Fourier component is proportional to the certainty (see main text Eq. (3)). This is trivially fulfilled for the cosine-shaped tuning curves that we used for illustration in the main text (main text Fig. 2). Here, we will demonstrate that this also holds for a more elaborate bump encoding scheme: specifically, we consider the case of a linear probabilistic population code (lPPC) (6–8) with independent Poisson neural noise. The central idea behind such an lPPC is that neuronal activity encodes an exponential family probability distribution, e.g., about HD, such that the natural parameters of this distribution can be retrieved through linear operations, that is, a weighted sum of neural activity.

In what follows, we will first show that an lPPC for a von Mises distribution with independent Poisson neurons gives rise to von Mises shape tuning curves, which are scaled by the encoded certainty (following (6)). Using this result, we will derive the population activity profile as a function of the encoded *estimate* and certainty that results from this encoding scheme, and show that the amplitude of this profile is indeed also proportional to the encoded certainty.

**A. Tuning with respect to (true) HD $\phi_t$.** We assume that tuning curves of the population encoding the posterior $p(\phi_t|V_t, Z_t)$ can be described by a typical shape $\tilde{f}$, which is scaled by the population gain $g$. That is, the tuning curve of a single neuron $i$ is given by $f_i(\phi_t) = g\,\tilde{f}_i(\phi_t)$. Following (6), we further assume that the neuronal population consists of $N$ independent Poisson neurons, which densely tile the stimulus space of true HDs, $\phi$. Thus, we can write down the probability of a population firing pattern $\mathbf{r} \in \mathbb{R}_+^N$ as

$$\begin{aligned} p(\mathbf{r}|\phi_t, g) &= \prod_i \frac{(g\tilde{f}_i(\phi_t))^{r_i}}{r_i!} \exp\left(-g\tilde{f}_i(\phi_t)\right) \\ &= \exp\left(\sum_i r_i \log(g\,\tilde{f}_i(\phi_t)) - \sum_i \log r_i! - \sum_i g\,\tilde{f}_i(\phi_t)\right) \\ &\propto_{\phi_t} \exp\left(\sum_i r_i \log \tilde{f}_i(\phi_t)\right), \end{aligned} \tag{S33}$$

where we used that $\sum_i g\,\tilde{f}_i(\phi_t)$ is approximately independent of HD $\phi_t$ due to the dense-tiling assumption.

Assuming that $p(\phi_t|\mathbf{r})$ follows an exponential family distribution, such as the von Mises distribution, an lPPC requires that the natural parameters of this distribution can be recovered from the population activity by a linear operation, i.e., a weighted sum. For a general exponential family distribution with $d$ sufficient statistics $\mathbf{T}(\phi_t) \in \mathbb{R}^d$, and natural parameters $\boldsymbol{\theta}$, we thus can re-parametrize the distribution in terms of the the population activities $\mathbf{r}$ (6):

$$\begin{aligned} p(\phi_t|\mathbf{r}) &= \frac{1}{Z(\phi_t, \boldsymbol{\theta})} \exp\left(\mathbf{T}(\phi_t)^T \cdot \boldsymbol{\theta}\right) \\ &= \frac{1}{Z(\phi_t, \mathbf{r})} \exp\left(\mathbf{T}(\phi_t)^T \cdot A\mathbf{r}\right), \end{aligned} \tag{S34}$$

where the decoder matrix $A \in \mathbb{R}^{d\times N}$ is defined via $\boldsymbol{\theta} = A\mathbf{r}$. Assuming a uniform prior over HD, that is, $p(\phi_t) \propto 1$, we can relate Eqs. [S33] and [S34] by Bayes' rule, $p(\phi_t|\mathbf{r}) \propto p(\mathbf{r}|\phi_t, g)$. This results in the following conditions for the tuning curves:

$$p(\phi_t|\mathbf{r}) \propto_{\phi_t} p(\mathbf{r}|\phi_t), \tag{S35}$$

$$\Rightarrow \log \tilde{\mathbf{f}}(\phi_t) = A^T \cdot \mathbf{T}(\phi_t). \tag{S36}$$

For a von Mises distribution, the natural parameters are given by $\mathbf{T}(\phi_t) = (\cos\phi_t, \sin\phi_t)^T$. Thus, the argument of the exponential in the neurons' tuning curves is a linear combination of sines and cosines. This, in turn, can be written as a single cosine $\propto c\cos(\phi_t - \phi_i)$, where $\phi_i \in [-\pi, \pi]$ denotes the "preferred HD" of neuron $i$. The tuning curve of a single neuron is thus von-Mises shaped, i.e.,

$$\tilde{f}_i(\phi_t) = \exp\left(\xi\cos(\phi_t - \phi_i)\right), \tag{S37}$$

where $\xi$ is an additional parameter that controls the width of the tuning curves. Furthermore, the decoder matrix is constrained via $(A^T)_i = \xi\,(\cos\phi_i, \sin\phi_i)$.

In order to determine the population gain $g$, note that we require the natural parameters of the von Mises distribution, $\boldsymbol{\theta} = \kappa_t\,(\sin\mu_t, \cos\mu_t)$, to be linearly decodable from the population activity via $\boldsymbol{\theta} = A\mathbf{r}$. Since $\boldsymbol{\theta}$ is proportional in $\kappa_t$, this linearity implies that the overall population activity $\mathbf{r}$ should also be overall scaled by $\kappa_t$. Hence, the tuning curve of a neuron with preferred HD $\phi_i$ reads:

$$f_i(\phi_t) = g\tilde{f}_i(\phi_t) = \kappa_t \exp\left(\xi\cos(\phi - \phi_i)\right). \tag{S38}$$

To summarize, an lPPC with independent Poisson neurons gives rise to von Mises shaped tuning curves, whose gain is scaled by the encoded certainty $\kappa_t$. Importantly, unlike for the encoded von Mises distribution, an increase in certainty $\kappa_t$ does not cause the resulting activity profile to sharpen.

**B. Tuning with respect to HD estimate $\mu_t$.** Tuning to true HD $\phi_t$ can only be measured if we have access to the encoded HD estimate. To instead find the tuning with respect to $\mu_t$ and $\kappa_t$ that parametrize the *distribution* of $\phi_t$, we need to average the neuron's tuning for a given $\mu_t$ and $\kappa_t$ over all possible realizations of $\phi_t$. This results in the following tuning with respect to $\mu_t$ and $\kappa_t$:

$$\begin{aligned} f_i(\mu_t, \kappa_t) &= \int_{-\pi}^{\pi} d\phi_t \, f_i(\phi_t) \, \mathcal{VM}\left(\phi_t; \mu_t, \kappa_t\right) \\ &= \frac{\kappa_t}{2\pi I_0(\kappa_t)} \int_{-\pi}^{\pi} d\phi_t \, \exp\left(\xi \cos(\phi_t - \phi_i) + \kappa_t \cos(\phi_t - \mu_t)\right) \\ &= \frac{\kappa_t}{2\pi I_0(\kappa_t)} \int_{-\pi}^{\pi} d\phi_t \, \exp\left(\tilde{\kappa}_{t,i} \cos(\phi_t - \tilde{\mu}_i)\right) \\ &= \kappa_t \frac{I_0(\tilde{\kappa}_{t,i})}{I_0(\kappa_t)}, \end{aligned} \qquad \text{[S39]}$$

with $\tilde{\kappa}_{t,i} = \sqrt{\xi^2 + \kappa_t^2 + 2\xi\kappa_t \cos(\phi_i - \mu_t)}$. This tuning curve is again bump-shaped, with a peak at the encoded HD estimate $\mu_t$ and the bump amplitude modulated by encoded certainty $\kappa_t$ in a nonlinear manner.

For small values of encoded certainty, the tuning curve approaches a cosine-shaped tuning with a gain that is a nonlinear function of $\kappa_t$. To see this, we use the series expansion of the Bessel function for a small argument $z$,

$$I_0(z) = \sum_{m=0}^{\infty} \frac{1}{m!\Gamma(m+1)} \left(\frac{z}{2}\right)^{2m} \approx 1 + \frac{1}{4} z^2 + \mathcal{O}(z^4), \qquad \text{[S40]}$$

and write for the tuning curve in the small-$\kappa_t$ limit

$$\kappa_t \frac{I_0(\tilde{\kappa}_{t,i})}{I_0(\kappa_t)} \approx \frac{\kappa_t}{I_0(\kappa_t)} \left(1 + \frac{1}{2}\tilde{\kappa}_{t,i}^2\right) = \frac{\kappa_t}{I_0(\kappa_t)} \left(1 + \frac{1}{4}\left(\xi^2 + \kappa_t^2 + \xi\kappa_t \cos(\phi_i - \mu_t)\right)\right). \qquad \text{[S41]}$$

Thus, the tuning curve of a neuron $i$ for small values of $\kappa_t$ is cosine-shaped, and modulated by the nonlinear factor $\frac{\xi\kappa_t^2}{4I_0(\kappa_t)}$, which asymptotically approaches $\frac{\xi}{4}\kappa_t^2$ for $\kappa_t \to 0$.

For large values of $\kappa_t$, the tuning curve is von-Mises shaped and the gain is asymptotically linear in encoded certainty. To see this, we use the Hankel expansion of the Bessel function $I_0(z)$ in the limit of large arguments $z$:

$$I_0(z) \approx \frac{e^z}{\sqrt{2\pi z z}} + \mathcal{O}\left(\frac{1}{z^2}\right), \qquad \text{[S42]}$$

and simplify

$$\kappa_t \frac{I_0(\tilde{\kappa}_{t,i})}{I_0(\kappa_t)} \approx \kappa_t \sqrt{\frac{\kappa_t}{\tilde{\kappa}_{t,i}}} \exp\left(\tilde{\kappa}_{t,i} - \kappa_t\right). \qquad \text{[S43]}$$

Taylor-expanding the exponent $\tilde{\kappa}_{t,i} - \kappa_t$ for small values of $1/\kappa_t$ yields,

$$\tilde{\kappa}_{t,i} - \kappa_t = \kappa_t \sqrt{1 + \frac{\xi^2}{\kappa_t^2} + \frac{\xi}{\kappa_t}\cos(\phi_i - \mu_t)} - \kappa_t \approx \frac{\xi}{2}\cos(\phi_i - \mu_t) + \frac{\xi^2}{2\kappa_t} + \mathcal{O}\left(\frac{1}{\kappa_t^2}\right). \qquad \text{[S44]}$$

Further, the pre-factor $\sqrt{\frac{\tilde{\kappa}_{t,i}}{\kappa_t}} \to 1$, and thus the tuning curve in the large-$\kappa_t$ limit reads:

$$f_i(\mu_t, \kappa_t) \to \kappa_t \exp\left(\frac{\xi}{2}\cos(\phi_i - \mu_t)\right). \qquad \text{[S45]}$$

The choice of the width parameter $\xi$ determines how large $\kappa_t$ has to be for the tuning curve to scale linearly with encoded certainty.

In Fig. S1, we demonstrate these limits (assuming $\xi = 1$ without loss of generality), and find numerically that linear scaling of the population activity amplitude holds well even for small $\kappa_t$ (e.g., $\kappa_t \sim 1$, cf. Fig. S1f). In addition, the width of the profile saturates quickly as we increase $\kappa_t$ (which indicates the transition from cosine-shaped to von-Mises shaped tuning curve), which makes the shape almost independent of $\kappa_t$. Therefore, the population profile is not just a rescaled version of the encoded probability distribution (Fig. S1c), because an increase in certainty does not cause the bump to sharpen indefinitely.

The linear scaling of the amplitude with $\kappa_t$, and (almost) constant width, indicate that the parameters of the von Mises distribution, $\mu_t$ and $\kappa_t$, can be retrieved from the population activity by computing the first Fourier coefficients:

$$\mathcal{F}_1^{\text{even}}\left[f_i(\mu_t, \kappa_t)\right] := \frac{1}{\pi}\int_{-\pi}^{\pi} d\phi_i \, f_i(\mu_t, \kappa_t)\cos(\phi_i) \propto \kappa_t \cos\mu_t = \theta_{t,1}, \qquad \text{[S46]}$$

$$\mathcal{F}_1^{\text{odd}}\left[f_i(\mu_t, \kappa_t)\right] := \frac{1}{\pi}\int_{-\pi}^{\pi} d\phi_i \, f_i(\mu_t, \kappa_t)\sin(\phi_i) \propto \kappa_t \sin\mu_t = \theta_{t,2}. \qquad \text{[S47]}$$

The certainty $\kappa_t$ can be retrieved via $\kappa_t = \sqrt{\theta_{t,1}^2 + \theta_{t,2}^2}$, and thus is proportional to the amplitude $c_1$ of the first Fourier component in amplitude-phase form. Likewise, the mean $\mu_t$ is the angle of the first Fourier component, i.e. $\mu_t = \arctan 2(\theta_{t,1}, \theta_{t,2})$. In other words, the tuning profile can be expanded as

$$f_i(\mu_t, \kappa_t) \sim \kappa_t \cos(\mu_t - \phi_i) + \mathcal{R}, \quad \text{[S48]}$$

where $\mathcal{R}$ collectively denotes the orthogonal other Fourier modes. In Fig. S1g-j, we confirm the proportionality of the amplitudes of the first Fourier coefficient in $\kappa_t$ numerically.

#### 3. Details on Bayesian ring attractor dynamics and parameter tuning

In the main text we consider a rate-based network model, called the *Bayesian ring attractor*, that implements an approximation to the circKF in the dynamics of its bump position and amplitude. Here, we derive this network in two steps. First, we start with a network that implements the circKF exactly (in the limit of an infinite number of neurons) by implementing the dynamics described by Eqs. [S19] and [S20]. This network won't be a ring attractor, as its activity will decay to zero in the absence of external inputs. After that we will change the network to instead implement the quadratic approximation to the circKF by implementing the dynamics described by Eqs. [S19] and [S21], resulting in the Bayesian ring attractor described in the main text.

Our derivation starts with a general network in the limit of infinitely many neurons, continuously covering the space of preferred HDs. For this network we will analytically derive dynamics of bump position and amplitude. Matching these dynamics to that of the circKF equations then allows us to determine the network parameters required for this implementation. The network we present in the main text is formulated for a finite number of neurons, and here we will further demonstrate that it is straightforward to change between those two representations. In fact, any network coefficients for the infinite-neuron network are chosen such that they also describe those used for the finite-neuron network in the main text.

**A. Network that exactly implements the circKF.** Let us make an ansatz for a continuous-space, linear network dynamics with an additional non-linear interaction term:

$$dr_t(\phi) = -\frac{1}{\tau} r_t(\phi)dt + g\left(r_t(\phi)\right) \cdot r_t(\phi)dt + \left(W * r_t\right)(\phi)\, dt + I_t^{\mathrm{ext}}(\phi). \qquad \text{[S49]}$$

Here, $r_t(\phi)$ denotes the activity of a neuron identified by its preferred HD $\phi$ at time $t$, and $I_t^{\mathrm{ext}}(\phi)$ is an external input. Due to the circular symmetry, the recurrent connectivity function $W(\Delta\phi)$ only depends on the relative distance $\Delta\phi$ between two neurons' preferred HD. Further, $\left(W * r_t\right)(\phi) := \frac{1}{\pi}\int d\phi' W(\phi-\phi') r_t(\phi)$ denotes a convolution.

We consider the decomposition of the activity profile $r_t(\phi)$ in terms of its Fourier modes:

$$r_t(\phi) = \frac{1}{2} r_0(t) + \sum_{k=1}^{\infty} \left( r_k^{\mathrm{even}}(t) \cos k\phi + r_k^{\mathrm{odd}}(t) \sin k\phi \right) \qquad \text{[S50]}$$

$$= \frac{1}{2} r_0(t) + \sum_{k=1}^{\infty} \tilde{r}_k(t) \cos k(\phi - \Psi_k(t)). \qquad \text{[S51]}$$

Note, that the Fourier coefficients $r_k^{\mathrm{even}}(t)$ and $r_k^{\mathrm{odd}}(t)$ are related to the coefficient's amplitude $\tilde{r}_k(t)$ and phase $\Psi_k(t)$ via a Cartesian to polar coordinate transformation. Taking the derivative on both sides (in the amplitude-phase form) results in:

$$dr_t(\phi) = \frac{1}{2}\, dr_0(t) + \sum_{k=1}^{\infty} \Big( \cos k(\phi - \Psi_k(t))\, d\tilde{r}_k(t) + k\tilde{r}_k(t) \sin k(\phi - \Psi_k(t))\, d\Psi_k(t) \Big). \qquad \text{[S52]}$$

Thus, we can determine the dynamics of the Fourier coefficients $r_0$, $\tilde{r}_k$, and $\Psi_k$ by Fourier-transforming Eq. [S49], and subsequently matching the coefficients in the Fourier modes:

$$dr_0(t) = \frac{1}{\pi}\int_{-\pi}^{\pi} d\phi\, (dr_t) = \left(-\frac{1}{\tau} + w_0\right) r_0(t)\, dt - g(r_t)\tilde{r}_0(t)\, dt + I_0^{\mathrm{ext}}(t), \qquad \text{[S53]}$$

$$\begin{aligned} d\tilde{r}_k(t) &= \frac{1}{\pi}\int_{-\pi}^{\pi} d\phi \cos k(\phi - \Psi_k(t))\, (dr_t) \\ &= \left(-\frac{1}{\tau} + w_k^{even}\right) \tilde{r}_k(t)\, dt - g(r_t)\tilde{r}_k(t)\, dt + I_k(t)\cos(\Phi_k(t) - \Psi_k(t)) \end{aligned} \qquad \text{[S54]}$$

$$\begin{aligned} d\Psi_k(t) &= \frac{1}{k\tilde{r}_k(t)} \frac{1}{\pi}\int_{-\pi}^{\pi} d\phi \sin k(\phi - \Psi_k(t))\, (dr_t) \\ &= \frac{w_k^{odd}}{k}\, dt + \frac{I_k(t)}{k\tilde{r}_k(t)} \sin(\Phi_k(t) - \Psi_k(t)), \end{aligned} \qquad \text{[S55]}$$

where we used the Fourier decompositions $W(\Delta\phi) = \frac{w_0}{2} + \sum_{k=1}^{\infty}\left(w_k^{\mathrm{even}}\cos(k\Delta\phi) + w_k^{\mathrm{odd}}\sin(k\Delta\phi)\right)$ and $I_t^{\mathrm{ext}}(\phi) = \frac{I_0}{2} + \sum_{k=1}^{\infty} I_k \cos(k(\phi - \Phi_k))$. Note that here, $I_k$ refers to the $k$-th Fourier amplitude of the input, and not to the modified Bessel function. Setting $\Psi_1(t) = \mu_t$ and $\tilde{r}_1(t) = \kappa_t$, the dynamics of the first Fourier components in amplitude-phase form read:

$$d\mu_t = w_1^{odd}\, dt + I_1(t)\sin(\Phi_1(t) - \mu_t), \qquad \text{[S56]}$$

$$d\kappa_t = \left(-\frac{1}{\tau} + w_1^{even}\right)\kappa_t\, dt - g(r_t)\kappa_t\, dt + I_1(t)\cos(\Phi_1(t) - \mu_t) \qquad \text{[S57]}$$

Comparing Eq. [S19] ($\mu_t$ from circKF) with Eq. [S56] and Eq. [S20] ($\kappa_t$ from circKF) with Eq. [S57] allows us to determine conditions for network parameters and external input in Eq. [S49], such that the circKF is exactly implemented in the dynamics of the network's first Fourier mode:

$$\begin{aligned} \text{Even recurrent connections} \quad & w_1^{\text{even}} = 1/\tau, \\ \text{Odd recurrent connections} \quad & w_1^{\text{odd}} = \frac{\kappa_v}{\kappa_\phi + \kappa_v} v_t, \\ \text{External input strength} \quad & I_1 = \sqrt{2\kappa_z dt}, \\ \text{External input phase} \quad & \Phi_1(t) = z_t, \\ \text{Nonlinear inhibition} \quad & g(r_t) = \frac{f(\kappa_t(r_t))}{2(\kappa_\phi + \kappa_v)}. \end{aligned}$$

Here, $v_t$ denotes the (observed) angular velocity with reliability $\kappa_v$, and $z_t$ the HD observation with reliability $\kappa_z$. The nonlinear inhibition needs to be able to compute the amplitude $\kappa_t$ from the network activity $r_t(\phi)$. Note that this does not impose any conditions on network parameters which do not affect the first Fourier component dynamics, for instance, higher order recurrent interaction strengths $w_k$ with $k \neq 1$. These can in principle be chosen freely.[‡] Note that, in this simple network, angular velocity observations modulate the first odd component of the recurrent connectivity matrix. This is biologically unrealistic, and will be addressed once we move to the multi-population network further below.

To summarize, one potential (out of many possible) network dynamics that implements the circKF in the dynamics of its first Fourier components reads:

$$dr_t(\phi) = -\frac{1}{\tau} r_t(\phi)\, dt - \frac{f(\kappa_t(r_t))}{2(\kappa_\phi + \kappa_v)} r_t(\phi) dt + \frac{1}{\tau} (\cos * r_t)(\phi) dt + \frac{\kappa_v}{\kappa_\phi + \kappa_v} v_t (\sin * r_t)(\phi)\, dt + I_t^{\text{ext}}(\phi). \qquad \text{[S58]}$$

Please consult Sec. D for an additional term required to account for $r_t$ being a stochastic process. We have not included this term here, as it only becomes important in the $dt \to 0$ limit, and does not contribute additional intuition about the network's operation.

**B. Network with quadratic nonlinearity.** While the network we have derived so far implements the cricKF exactly, its activity decays to zero in the absence of external inputs, such that it is not an attractor network. In this section we will instead use the quadratic approximation to the circKF, which will lead to the Bayesian ring attractor we discuss in the main text. To do so, we use the following nonlinearity for the inhibitory interaction:

$$g(r_t) r_t = w^{\text{quad}} (M * [r_t]_+)(\phi) \circ r_t(\phi), \qquad \text{[S59]}$$

with rectification nonlinearity $[\cdot]_+$ and a constant function $M = \frac{\pi}{2}$. Here, $\circ$ denotes the Hadamar (piecewise) product. In the main text, we wrote this interaction as $g(r_t) r_t \to w^{\text{quad}} \left(\pi \sum_{i=1}^N [r_t^{(i)}]_+\right) \cdot r_t$, which is equivalent, but less technical.

We assume $r_t$ to be dominated by its first Fourier component, such that the other orders become negligible, i.e. $r_t(\phi) = \kappa_t \cos(\phi - \mu_t) + \mathcal{R}$ with $\mathcal{R}$ small.[§] We find

$$(M * [r_t]_+)(\phi) \approx \frac{1}{\pi} \int_{-\pi}^{\pi} d\phi' \frac{\pi}{2} \left[\kappa_t \cos(\phi' - \mu_t)\right]_+ = \kappa_t. \qquad \text{[S60]}$$

Fourier-transforming the nonlinearity with respect to the amplitude-phase form yields:

$$\begin{aligned} \frac{1}{\pi} \int_{-\pi}^{\pi} d\phi' \cos(\phi' - \mu_t) g(r_t) r_t &= \frac{w^{\text{quad}}}{\pi} \int_{-\pi}^{\pi} d\phi' \, \cos(\phi' - \mu_t) \left(M * [r_t]_+\right)(\phi') \cdot r_t(\phi') \\ &= \frac{w^{\text{quad}}}{\pi} \kappa_t \int_{-\pi}^{\pi} d\phi' \, \cos(\phi' - \mu_t) \, r_t(\phi') = w^{\text{quad}} \kappa_t^2. \end{aligned} \qquad \text{[S61]}$$

Thus, the dynamics of the first Fourier amplitude of a network with this nonlinearity is given by:

$$d\kappa_t = \left(-\frac{1}{\tau} + w_1^{even}\right) \kappa_t \, dt - w^{\text{quad}} \kappa_t^2 \, dt + I_1(t) \cos(\Phi_1(t) - \mu_t). \qquad \text{[S62]}$$

The network parameters can be tuned such that the dynamics match that of the quadratic approximation of the circular Kalman filter (Eq. [S19] and [S21]), analogously to the previous section. This yields the following network parameters for a Bayesian ring-attractor network:

[‡] Practically, we chose them such that higher-order Fourier modes and the zero-th mode decay reasonably fast, to produce a unimodal activity bump.

[§] Alternatively, we can consider additionally convolving $r_t$ with a cosine before applying the rectification, effectively filtering out the desired mode.

$$
\begin{aligned}
\text{Even recurrent connections} \quad & w_1^{\text{even}} = 1/\tau + \frac{1}{\kappa_\phi + \kappa_v}, \\
\text{Odd recurrent connections} \quad & w_1^{\text{odd}} = \frac{\kappa_v}{\kappa_\phi + \kappa_v} v_t, \\
\text{External input strength} \quad & I_1 = \sqrt{2\kappa_z dt}, \\
\text{External input phase} \quad & \Phi_1(t) = z_t, \\
\text{Quadratic inhibition} \quad & w^{\text{quad}} = \frac{1}{\kappa_\phi + \kappa_v},
\end{aligned}
$$

**C. Continuous vs. discrete networks.** The analysis we have presented above is valid for a continuum of neurons, i.e. $N \to \infty$, that span a continuum of preferred HDs. Formally, this implies that the difference in preferred HD between two 'neighboring' neurons converges to zero, $\Delta\phi := \phi_i - \phi_j = \frac{2\pi}{N} \to 0$. In the text and for our simulations, we used a discretized network, where we assumed the preferred HDs of the neurons to be equally spaced, but finite.

It is straightforward to go back and forth between these two representations (cf. (9)): in a discretized network, $\mathbf{r}_t$ denotes a vector of neural activities, indexed by their preferred HD $\phi_i$, which becomes a function $r_t(\phi)$ for a continuous network. Likewise, connectivity matrices $W$ become functions with two arguments $W(\phi_i, \phi_j)$, and matrix multiplications become integrals. The circular symmetry of HD implies that the entries of a connectivity matrix only depend on the relative distance between two neurons, and not on absolute position, such that for a connectivity matrix $W$ we can write $W_{ij} = W(\phi_i, \phi_j) = W(\phi_i - \phi_j)$. Thus, we can write matrix multiplications as convolutions (assuming the vectors and matrix are ordered with respect to their preferred HD):

$$
(W \cdot \mathbf{r}_t)_i = \sum_{j=1}^{N} W_{ij} r_{t,j} = \frac{N}{2\pi} \sum_{j=1}^{N} W_{ij} r_{t,j} \Delta\phi \tag{S63}
$$

$$
\overset{N\to\infty, \Delta\phi\to 0}{\to} \frac{N}{2\pi} \int_{-\pi}^{\pi} d\phi' W(\phi, \phi') r_t(\phi') = \frac{N}{2\pi} \int_{-\pi}^{\pi} d\phi' W(\phi - \phi') r_t(\phi') = \frac{N}{2} \left(W * r_t\right)(\phi). \tag{S64}
$$

where we defined the convolution as above. Thus, to ensure consistency between the coefficients of the matrices used in the main text and the coefficients of the connectivity functions we used in our analysis in the SI, we scaled the connectivity matrices in the main text by a factor $\frac{2}{N}$.

**D. Stochastic correction [Technical].** The derivation in the previous section did not take into account that due to the dependence on the angular velocity observations $v_t$, the phase $\Psi_k(t)$ is actually an Itô stochastic process, and hence the network activity $r_t$ is, too. Thus, when performing a change of variables, such as the expansion Eq. [S52], we have to use Itô's lemma (Eq. [S24]), and expand up to second order in $\Psi_k(t)$ (we have seen that the dynamics of the amplitude $\tilde{r}_k(t)$ is independent of $v_t$, and thus only carries first order terms):

$$
dr_t(\phi) = d\left(\frac{1}{2} r_0(t) + \sum_{k=1}^{\infty} \tilde{r}_k(t) \cos k(\phi - \Psi_k(t))\right) \tag{S65}
$$

$$
\begin{aligned}
&= \frac{1}{2} dr_0(t) + \sum_{k=1}^{\infty} \Big( \cos k(\phi - \Psi_k(t)) d\tilde{r}_k(t) + k \tilde{r}_k(t) \sin k(\phi - \Psi_k(t)) d\Psi_k(t) \\
&\qquad - \frac{1}{2} k^2 \tilde{r}_k(t) \cos k(\phi - \Psi_k(t)) (d\Psi_k(t))^2 \Big),
\end{aligned} \tag{S66}
$$

This implies that, if we take the effect of stochastic processes into account, comparing the Fourier coefficients in amplitude-phase form will not single out the dynamics of the amplitude $d\tilde{r}_k$, because there are now two terms proportional to $\cos k(\phi - \Psi_k(t))$. Fortunately, the problem can be solved "backwards" using the analogy to coordinate transforms in Section C, thereby restricting ourselves to the first Fourier mode (higher modes are analogous): First, we perform the Fourier transform of the dynamics in Cartesian coordinates, i.e., with respect to $\cos(\phi)$ and $\sin(\phi)$. We then note that changing this into amplitude-phase form is mathematically equivalent to a coordinate transform between the natural parameters of the von Mises distribution and the $\mu$,$\kappa$-parametrization. Next, we require that such a coordinate transform ought to result in the dynamics for $\mu_t$ and $\kappa_t$ in Eq. [S56] and [S57]. Using the analogy to Section C, we find that an additional decay term $-\frac{1}{2} \frac{\kappa_v/\kappa_\phi}{\kappa_v + \kappa_\phi} r_t(\phi) dt$ is needed in the network dynamics, which implements the Itô correction on the level of the natural parameters (cf. Eq. [S26]). Apart from this additional decay, the conditions on the other network parameters remains unchanged.

This stochastic correction is not strictly needed to gain intuition about the theory, and if anything, the use of continuous-time stochastic calculus seems to make things *less* intuitive. Practically, we used an additional decay term in Eq. [S58] whenever the angular velocity observations were drawn from the true generative model and the time step $dt$ was small enough to justify the notion of "continuous time", which was the case for all our simulations.

#### 4. Details on *Drosophila*-like network

Relying on large-scale connectomics data of the *Drosophila* HD system (10, 11), we now ask if a Bayesian ring attractor can be implemented in a network that obeys biological network connectivity constraints. Here we show how the motifs of this network – and, by extension, any biological ring attractor network – could potentially implement dynamic Bayesian inference.

**A. Connectivity motifs in the *Drosophila* HD system connectome.** The ring attractor in the *Drosophila* HD system is composed of three core cell types, called EPG, PEN1 and Δ7 neurons (10–12), cf. Fig. S4a-c. HD is represented as a bump of neural activity in the EPG population (13). These neurons are recurrently connected with excitatory PEN1 neurons. When the fly turns, this differentially activates PEN1 neurons in the right and left brain hemispheres, and because PEN1 neurons have asymmetric (shifted) projections back to EPG neurons, they can rotate the bump of EPG activity in accordance with the fly's rotation (14, 15). This motif effectively establishes the velocity-modulated odd recurrent connectivity required to initiate turns in ring attractor networks (Fig. S4d). Moreover, EPG neurons are recurrently connected with inhibitory Δ7 neurons, which establishes broad inhibition (Fig. S4e). Finally, EPG neurons receive inhibitory inputs from so-called ER neurons, which send HD information to EPG neurons (16–18) (Fig. S4f). In summary, the fly's HD system is equipped with the basic motifs to implement a Bayesian ring attractor.

**B. A multi-network model mimicking the *Drosophila* HD system connectome.** The main idea of the idealized network in the previous section was to tune the network parameters such that the circKF (or the quadratic approximation of the circKF) was implemented in the coefficients of the first Fourier mode. Here, we will use the connectome of the fruit fly *Drosophila* (10) to build a recurrent neural network, and show that the quadratic approximation of the circKF can be implemented in such an architecture by determining the coefficients analogously. Thereby, we first approximate the connectivity matrices describing this connectome (Fig. S4b) by analytically accessible functions, which nonetheless retain the main features of this connectivity (as outlined, e.g., in (12)), and preserve the motifs that implement the ring-attractor in the *Drosophila* HD system (see review in (19)). We in turn analytically determine the conditions for the coefficients of the connectivities between (rather than within) the different network populations, such that the dynamics of the first Fourier components match that of the quadratic approximation of the circKF.

Specifically, we consider five neuronal populations: an HD population, $r^{HD}$, which we designed to track HD estimate and certainty with its bump parameter dynamics, two angular ($\mathrm{AV}^+$ and $\mathrm{AV}^-$) velocity populations, $r^{AV^+}$ and $r^{AV^-}$, which are tuned to head direction and are differentially modulated by angular velocity input, an inhibitory (INH) population, $r^{INH}$, and a population $I^{ext}$ that represents external input, that is, the HD observations. As before, the population activities $r(\phi)$ are functions of preferred HDs, $\phi$, but we will drop the argument $\phi$ to keep the notation uncluttered.

We start with the following ansatz for a network dynamics:

$$\begin{aligned} dr_t^{HD} =& -\frac{1}{\tau_{HD}} r_t^{HD}\,dt + W_{HD\leftarrow HD} * r_t^{HD}\,dt + W_{HD\leftarrow AV^+} * r_t^{AV^+} + W_{HD\leftarrow AV^-} * r_t^{AV^-}\,dt \\ &+ \left(W_{HD\leftarrow INH} * [r_t^{INH}]_+\right) \circ r_t^{HD}\,dt + I_t^{ext}, \end{aligned} \quad [S67]$$

$$dr_t^{AV^+} = \frac{1}{\tau_{AV^+}}\left(-r_t^{AV^+} + (o^{AV} + v_t) W_{AV^+\leftarrow HD} * r_t^{HD}\right) dt, \quad [S68]$$

$$dr_t^{AV^-} = \frac{1}{\tau_{AV^-}}\left(-r_t^{AV^-} + (o^{AV} - v_t) W_{AV^-\leftarrow HD} * r_t^{HD}\right) dt, \quad [S69]$$

$$dr_t^{INH} = \frac{1}{\tau_{INH}}\left(-r_t^{INH} + W_{INH\leftarrow HD} * [r_t^{HD}]_+ + W_{INH\leftarrow INH} * r_t^{INH}\right) dt. \quad [S70]$$

From the connectivity profile ((10), cf. Fig. S4b), we make the following ansatz for the connectivity functions (which results in Fig. S4c):

$$W_{HD\leftarrow HD}(\Delta\phi) = c_0^{HD} + c_1^{HD}\left[\cos\Delta\phi\right], \quad [S71]$$

$$W_{AV^\pm\leftarrow HD}(\Delta\phi) = c^{AV^\pm\leftarrow HD}\delta(\Delta\phi), \quad [S72]$$

$$W_{HD\leftarrow AV^\pm}(\Delta\phi) = c^{HD\leftarrow AV^\pm}\left[\sin\left(\Delta\phi \pm \frac{\pi}{4}\right)\right]_+, \quad [S73]$$

$$W_{INH\leftarrow HD}(\Delta\phi) = \frac{c_0^{INH\leftarrow HD}}{2} + c_1^{INH\leftarrow HD}\cos(\Delta\phi), \quad [S74]$$

$$W_{INH\leftarrow INH} = \frac{c_0^{INH\leftarrow INH}}{2} + c_1^{INH\leftarrow INH}\cos(\Delta\phi), \quad [S75]$$

$$W_{HD\leftarrow INH}(\Delta\phi) = c^{HD\leftarrow INH}\delta(\Delta\phi). \quad [S76]$$

In what follows, we will derive the conditions for the connection strengths in this ansatz that allow an implementation of the quadratic approximation of the circKF in the dynamics of the first Fourier component. Thereby, we make the assumption that the leading order of the HD population activity $r_t^{HD}$ is a cosine, i.e. $r_t^{HD}(\phi) = \frac{r_0^{HD}(t)}{2} + \kappa_t\cos(\phi - \mu_t) + \mathcal{R}$, and that higher-order Fourier modes are negligible. We further assume that the time constants of the $\mathrm{AV}^\pm$ and INH populations, $\tau_{AV^\pm}$ and $\tau_{INH}$, are much smaller than $\tau_{HD}$ of the HD population, which allows us to assume that the activity in those populations is stationary.

***B.1. AV$^\pm$ population.*** As described above, the integration of turning signals in the fruit fly is modulated through differential activation of PEN1 neurons (our AV$^\pm$ population) in the right and left brain hemispheres that asymmetrically project back to EPG neurons (our HD population) (14, 15). This motif implements the effective asymmetric angular velocity-dependent recurrent connectivity that is needed to rotate the activity in ring-attractor networks (20, 21). Thus, we will tune the parameters in the HD→AV$^\pm$ →HD circuit such that the resulting effective odd recurrent connectivity contribution $w_1^{odd}$ (i.e., that proportional to $\sin(\phi - \mu_t)$ implements the turn in the activity profile due to angular velocity integration, cf. Eq. [S55]).

As a first step, we compute the activities in the AV$^\pm$ populations. It is straightforward to check that, if the time constant $\tau_{AV} \ll \tau_{HD}$, the activity in the AV populations can be described by its stationary activity at every point in time:

$$\begin{aligned} r_t^{AV^\pm} &= (o_{AV} \pm v_t) W_{AV^\pm \leftarrow HD} * r_t^{HD} = c^{AV^\pm \leftarrow HD} (o_{AV} \pm v_t) \frac{1}{\pi} \int_{-\pi}^{\pi} d\phi' \delta(\phi - \phi') r_t^{HD}(\phi') \\ &= c^{AV^\pm \leftarrow HD} (o_{AV} \pm v_t)\, r_t^{HD}. \end{aligned} \tag{S77}$$

Expanding the connectivity function from the HD to the AV$^\pm$ populations in a Fourier series yields:

$$W_{HD \leftarrow AV^\pm} = c^{HD \leftarrow AV^\pm} \left[ \pm \sin(\Delta\phi \pm \frac{\pi}{4}) \right]_+ = c^{HD \leftarrow AV^\pm} \left( \frac{1}{\pi} + \frac{1}{2\sqrt{2}} \cos(\Delta\phi) \pm \frac{1}{2\sqrt{2}} \sin(\Delta\phi) \right) + \mathcal{R}, \tag{S78}$$

allowing us to compute the effective recurrent contributions in the HD population that is mediated via this network motif:

$$\begin{aligned} W_{HD \leftarrow AV^+} * r_t^{AV^+} &= c^{HD \leftarrow AV^\pm} c^{AV^\pm \leftarrow HD} (o_{AV} + v_t) \frac{1}{\pi} \int_{-\pi}^{\pi} d\phi' \left( \frac{1}{\pi} + \frac{1}{2\sqrt{2}} \cos(\phi - \phi') + \frac{1}{2\sqrt{2}} \sin(\phi - \phi') + \mathcal{R} \right) r_t^{HD}(\phi') \\ &= c^{HD \leftarrow AV^\pm} c^{AV^\pm \leftarrow HD} (o_{AV} + v_t) \left( \frac{r_0^{HD}}{\pi} + \frac{\kappa_t}{2\sqrt{2}} \cos(\phi - \mu_t) + \frac{\kappa_t}{2\sqrt{2}} \sin(\phi - \mu_t) \right), \end{aligned} \tag{S79}$$

$$W_{HD \leftarrow AV^-} * r_t^{AV^-} = c^{HD \leftarrow AV^\pm} c^{AV^\pm \leftarrow HD} (o_{AV} - v_t) \left( \frac{r_0^{HD}}{\pi} + \frac{\kappa_t}{2\sqrt{2}} \cos(\phi - \mu_t) - \frac{\kappa_t}{2\sqrt{2}} \sin(\phi - \mu_t) \right), \tag{S80}$$

and thus

$$\begin{aligned} W_{HD \leftarrow AV^+} * r_t^{AV^+} &+ W_{HD \leftarrow AV^-} * r_t^{AV^-} \\ &= c^{HD \leftarrow AV^\pm} c^{AV^\pm \leftarrow HD} \left( 2 \frac{o_{AV}}{\pi} r_0^{HD} + \kappa_t \frac{o_{AV}}{\sqrt{2}} \cos(\phi - \mu_t) + \kappa_t v_t \frac{1}{\sqrt{2}} \sin(\phi - \mu_t) \right). \end{aligned} \tag{S81}$$

Thus, this motif implements an effective odd recurrent connectivity with $w_1^{odd} = \frac{c^{HD \leftarrow AV^\pm} c^{AV^\pm \leftarrow HD}}{\sqrt{2}} v_t$. We require that the effective odd recurrent connectivity is the same as in the Bayesian ring attractor, that is,

$$w_1^{\text{odd}} = \frac{c^{HD \leftarrow AV^\pm} c^{AV^\pm \leftarrow HD}}{\sqrt{2}} v_t \overset{!}{=} \frac{\kappa_v}{\kappa_\phi + \kappa_v} v_t, \tag{S82}$$

and thus the condition for the coefficients reads:

$$c^{HD \leftarrow AV^\pm} = \frac{\sqrt{2}}{c^{AV^\pm \leftarrow HD}} \frac{\kappa_v}{\kappa_\phi + \kappa_v}. \tag{S83}$$

Interestingly, due to the offset $o_{AV}$ we also obtain a recurrent contribution to the activity baseline $r_0(t)$, and a contribution to the *even* first order recurrent connectivity,

$$w_1^{\text{even, AV}} = c^{HD \leftarrow AV^\pm} c^{AV^\pm \leftarrow HD} \frac{o_{AV}}{\sqrt{2}} \tag{S84}$$

$$= \frac{\kappa_v}{\kappa_\phi + \kappa_v} o_{AV}. \tag{S85}$$

We will return to this when computing the recurrent connectivities within the HD populations below.

***B.2. INH population..*** In our network, the recurrent interaction with the INH population implements the quadratic inhibition. In the same way we tracked the effective odd recurrent through the AV$^\pm$ recurrent loop, we will here determine the effective quadratic interaction strength $w^{\text{quad}}$ as a function of the network parameters, and then tune it in order to implement the quadratic approximation of the circKF.

To determine the activity in the INH population, we first expand $\left[r_t^{HD}\right]_+$ in its Fourier series:

$$\begin{aligned}\left[r_t^{HD}\right]_+ &\approx \left[\frac{r_0^{HD}}{2} + \kappa_t \cos(\phi - \mu_t)\right]_+ \\ &\approx \frac{r_0^{HD}\phi_c}{2\pi} + \frac{\kappa_t}{\pi}\sin\phi_c + \left(\frac{\kappa_t}{\pi}\phi_c + \frac{r_0^{HD}}{2\pi}\sin\phi_c\right)\cos(\phi-\mu_t) + \mathcal{R} \\ &\approx \frac{r_0^{HD}}{4} + \frac{\kappa_t}{\pi} + \left(\frac{\kappa_t}{2} + \frac{r_0^{HD}}{\pi}\right)\cos(\phi-\mu_t) + \mathcal{R},\end{aligned} \tag{S86}$$

with cutoff angle $\phi_c = \arccos\left(-\frac{\kappa_t}{2r_1^{HD}}\right) \approx \frac{\pi}{2} + \frac{r_0^{HD}}{2\kappa_t}$ for $\kappa_t \gg r_0^{HD}/2$. With the dynamics of the INH population in Eqs. [S70], and the connectivity functions in [S74] and [S75], we can write down the dynamics of the first two Fourier coefficients in the INH population:

$$\tau_{INH}\, dr_0^{INH} = \left(-r_0^{INH} + \left(\frac{r_0^{HD}}{2} + \frac{2}{\pi}\kappa_t\right)c_0^{INH\leftarrow HD} + c_0^{INH\leftarrow INH} r_0^{INH}\right)dt, \tag{S87}$$

$$\tau_{INH}\, dr_1^{INH} = \left(-r_1^{INH} + \left(\frac{1}{2}\kappa_t + \frac{1}{\pi}r_0^{HD}\right)c_1^{INH\leftarrow HD} + c_1^{INH\leftarrow INH} r_1^{INH}\right)dt. \tag{S88}$$

Assuming again that the dynamics in the INH population is much faster than in the HD population, $\tau_{INH} \ll \tau_{HD}$, we can write down the stationary activities of the activity profile in the INH population:

$$r_0^{INH} = \frac{\frac{r_0^{HD}}{2} + \frac{2}{\pi}\kappa_t}{1 - c_0^{INH\leftarrow INH}} c_0^{INH\leftarrow HD}, \tag{S89}$$

$$r_1^{INH} = \frac{\frac{1}{2}\kappa_t + \frac{1}{\pi}r_0^{HD}}{1 - c_1^{INH\leftarrow INH}} c_1^{INH\leftarrow HD}. \tag{S90}$$

Plugging this into Eq. [S67], we obtain the change in the amplitude of the first Fourier mode through the interaction with the INH population:

$$\begin{aligned}\left(W_{HD\leftarrow INH} * [r_t^{INH}]_+\right)\cdot r_t^{HD} &= c^{HD\leftarrow INH}\left(\frac{r_0^{INH}}{2} + r_1^{INH}\cos(\phi-\mu_t)\right)\cdot r_t^{HD}(\phi) \\ &= c^{HD\leftarrow INH}\left(\frac{r_0^{INH}}{2}\kappa_t + r_1^{INH}\frac{r_0^{HD}}{2}\right)\cos(\phi-\mu_t) + \mathcal{R} \\ &= c^{HD\leftarrow INH}\left[\frac{c_0^{INH\leftarrow HD}}{\pi(1-c_0^{INH\leftarrow INH})}\kappa_t^2 + \left(\frac{c_0^{INH\leftarrow HD}}{1-c_0^{INH\leftarrow INH}} + \frac{c_1^{INH\leftarrow HD}}{1-c_1^{INH\leftarrow INH}}\right)\frac{r_0^{HD}}{4}\kappa_t\right. \\ &\quad \left. + \frac{c_1^{INH\leftarrow HD}}{\pi(1-c_1^{INH\leftarrow INH})}\frac{(r_0^{HD})^2}{2}\right]\cos(\phi-\mu_t).\end{aligned} \tag{S91}$$

The first term on the right hand side has our desired quadratic interaction. It matches that of the quadratic approximation of the circKF $w^{quad} = 1/(\kappa_\phi + \kappa_v)$, if the following condition is fulfilled:

$$c^{HD\leftarrow INH} = -\frac{1}{\kappa_\phi + \kappa_v}\frac{\pi(1 - c_0^{INH})}{c_0^{INH\leftarrow HD}}. \tag{S92}$$

The other terms in Eq. [S91] are "nuisance" terms, which, if too large, may significantly interfere with the inference dynamics. However, if $r_0^{HD}$ is small compared to $\kappa_t$, which we confirmed in simulations to be generally the case, the effect of the nuisance terms is negligible. This can further be stabilized by choosing $|c_1^{INH\leftarrow HD}| \ll |1 - c_1^{INH\leftarrow INH}|$. Interestingly, this implies that certainty $\kappa_t$ mainly governs the activity in the *zero-th* order of the INH activity (Eq. [S89]).

***B.3. Recurrent excitation within HD population.*** In the same spirit as above, here we compute the effective even recurrent connectivity of the network in order to match it with recurrent interaction $w_1^{\text{even}}$ in the network implementation of the circKF. Starting from the Fourier expansion of the recurrent connectivity,

$$W_{HD\leftarrow HD} = c_0^{HD} + c_1^{HD}\left[\cos(\Delta\phi)\right]_+ \approx c_0^{HD} + \frac{c_1^{HD}}{\pi} + \frac{c_1^{HD}}{2}\cos(\Delta\phi) + \mathcal{R}, \tag{S93}$$

we determine the change in activity due to the recurrent interaction within the HD population:

$$W_{HD\leftarrow HD} * r_t^{HD} = \left(c_0^{HD} + \frac{c_1^{HD}}{\pi}\right)r_0^{HD} + \frac{c_1^{HD}}{2}\kappa_t\cos(\phi-\mu_t) + \mathcal{R}. \tag{S94}$$

Recall that the interaction with the AV$^{\pm}$ populations also induced an effective *even* recurrent connectivity (Eq. [S85]), such that the overall even recurrent connectivity in the network is given by,

$$w_1^{\text{even}} = w_1^{\text{even, HD}} + w_1^{\text{even, AV}} = \frac{c_1^{HD}}{2} + \frac{\kappa_v}{\kappa_\phi + \kappa_v} o_{AV} \stackrel{!}{=} \frac{1}{\tau} + \frac{1}{\kappa_\phi + \kappa_v}. \tag{S95}$$

This defines the following condition for the recurrent interaction within the HD population:

$$c_1^{HD} = 2\left(\frac{1}{\tau} + \frac{1}{\kappa_\phi + \kappa_v} - \frac{\kappa_v}{\kappa_\phi + \kappa_v} o_{AV}\right). \tag{S96}$$

The zero-order contribution in Eq. [S94] multiplying $c_1^{HD}$ is significant, and exceeds the first-order interaction in magnitude, which makes the network unstable. We thus require a negative constant recurrent connectivity to balance this zero-order contribution, chosen such that this contributions in the dynamics of $r_0^{HD}$ decays over time:

$$2\left(c_0^{HD} + \frac{c_1^{HD}}{\pi}\right) \stackrel{!}{<} \frac{1}{\tau}, \tag{S97}$$

and thus we arrive at our final condition:

$$c_0^{HD} < \frac{1}{2\tau} - \frac{c_1^{HD}}{\pi}. \tag{S98}$$

***B.4. Summary of network connectivities.*** To summarize, we analytically determined that the following connectivity matrices in the network dynamics in Eq. [S67]-[S70] implement the quadratic approximation of the circKF in the HD population. As a reminder, these network dynamics are:

$$\begin{aligned}
dr_t^{HD} &= -\frac{1}{\tau_{HD}} r_t^{HD}\, dt + W_{HD\leftarrow HD} * r_t^{HD}\, dt + W_{HD\leftarrow AV^+} * r_t^{AV^+} + W_{HD\leftarrow AV^-} * r_t^{AV^-}\, dt \\
&\quad + \left(W_{HD\leftarrow INH} * [r_t^{INH}]_+\right) \circ r_t^{HD}\, dt + I_t^{ext}, \\
dr_t^{AV^+} &= \frac{1}{\tau_{AV^+}}\left(-r_t^{AV^+} + (o^{AV} + v_t) W_{AV^+\leftarrow HD} * r_t^{HD}\right) dt \\
dr_t^{AV^-} &= \frac{1}{\tau_{AV^-}}\left(-r_t^{AV^-} + (o^{AV} - v_t) W_{AV^-\leftarrow HD} * r_t^{HD}\right) dt \\
dr_t^{INH} &= \frac{1}{\tau_{INH}}\left(-r_t^{INH} + W_{INH\leftarrow HD} * [r_t^{HD}]_+ + W_{INH\leftarrow INH} * r_t^{INH}\right) dt.
\end{aligned}$$

**Recurrent excitation within HD population:**

$$\begin{aligned}
(W_{HD\leftarrow HD})_{ij} &= \frac{2}{N_{HD}}\left(c_0^{HD} + c_1^{HD}\left[\cos\left(\phi_i^{HD} - \phi_j^{HD}\right)\right]_+\right), \\
\text{with}\ \ c_1^{HD} &= 2\left(\frac{1}{\kappa_\phi + \kappa_v} + \frac{1}{\tau_{HD}} - o^{AV}\frac{\kappa_v}{\kappa_\phi + \kappa_v}\right), \quad c_0^{HD} < \frac{1}{2\tau} - \frac{c_1^{HD}}{\pi}.
\end{aligned} \tag{S99}$$

**Recurrent excitation between HD and AV+ and AV- populations:**

$$(W_{AV^\pm\leftarrow HD})_{ij} = c^{AV^\pm\leftarrow HD}\delta_{ij}, \tag{S100}$$

$$\begin{aligned}
(W_{HD\leftarrow AV^\pm})_{ij} &= \frac{2}{N_{AV^\pm}} c^{HD\leftarrow AV^\pm}\left[\sin\left(\phi_i^{HD} - \phi_j^{AV^\pm} \pm \frac{\pi}{4}\right)\right]_+, \\
\text{with}\ \ c^{HD\leftarrow AV^\pm} &= \frac{\sqrt{2}}{c^{AV^\pm\leftarrow HD}} \frac{\kappa_v}{\kappa_\phi + \kappa_v}.
\end{aligned} \tag{S101}$$

**Recurrent inhibition between HD and INH populations:**

$$(W_{INH\leftarrow HD})_{ij} = \frac{2}{N_{HD}}\left(\frac{c_0^{INH\leftarrow HD}}{2} + c_1^{INH\leftarrow HD}\cos\left(\phi_i^{INH} - \phi_j^{HD}\right)\right), \qquad \text{[S102]}$$

$$(W_{INH\leftarrow INH})_{ij} = \frac{2}{N_{INH}}\left(\frac{c_0^{INH}}{2} + c_1^{INH}\cos\left(\phi_i^{INH} - \phi_j^{HD}\right)\right), \quad \text{with } |c_1^{INH\leftarrow HD}| << |1 - c_1^{INH}| \qquad \text{[S103]}$$

$$(W_{HD\leftarrow INH})_{ij} = c^{HD\leftarrow INH}\delta_{ij}, \quad \text{with } c^{HD\leftarrow INH} = -\frac{1}{\kappa_\phi + \kappa_v}\frac{\pi(1 - c_0^{INH})}{c_0^{INH\leftarrow HD}}. \qquad \text{[S104]}$$

Activities of the EXT population were assumed to give rise to a bump-shaped inhibitory input opposite of the HD observation, loosely related to how ring neurons mediate such input to the EPG neurons (17, 18). We thus modeled this bump-shaped input to the HD population directly without explicitly representing a dynamics of the EXT population.

**External input:**

$$I_{i,t}^{ext} = -2\sqrt{2\kappa_z dt}\left[\cos(\phi_i^{HD} - z_t + \pi)\right]_+. \qquad \text{[S105]}$$

The network dynamics still has a considerable number of degrees of freedom. That is, the baseline $o^{AV}$, network connectivity strengths $c^{AV^\pm\leftarrow HD}$, $c_0^{INH\leftarrow HD}$, $c_1^{INH\leftarrow HD}$, $c_0^{INH}$, $c_1^{INH}$, and time scales $\tau_{HD}$, $\tau_{AV+}$, $\tau_{AV-}$ and $\tau_{INH}$ can essentially be chosen freely. If the number of neurons $N$ differs between populations, the $\delta_{ij}$'s can be replaced by a normalized, Gaussian-shaped kernel with a finite width. For our analytical results to hold, we require $\tau_{HD} \gg \tau_{AV+}, \tau_{AV-}, \tau_{INH}$. We further constrained the network by choosing $c_0^{INH\leftarrow HD} > 0$, $c_1^{INH\leftarrow HD} \leq 0$ and $|c_0^{INH\leftarrow HD}| > |c_1^{INH\leftarrow HD}|$, which leads to the broad excitatory input into the INH population, and the formation of an 'antibump', similarly to the one observed in $\Delta 7$ neurons (12).

**C. Drosophila-like network HD tracking performance.** To demonstrate that the mutli-population network can indeed implement the quadratic approximation to the circKF, we measured its HD tracking performance and compared it to the circKF and the Bayesian ring attractor. As shown in Fig. S4e & f, this confirmed that this network indeed achieves a HD tracking performance indistinguishable to that of our idealized Bayesian ring attractor network. Thus, even when we add the constraints dictated by the actual connectivity patterns of neural networks in the brain, the resulting network is still able to implement dynamic Bayesian inference.

#### 5. Supplementary Figures

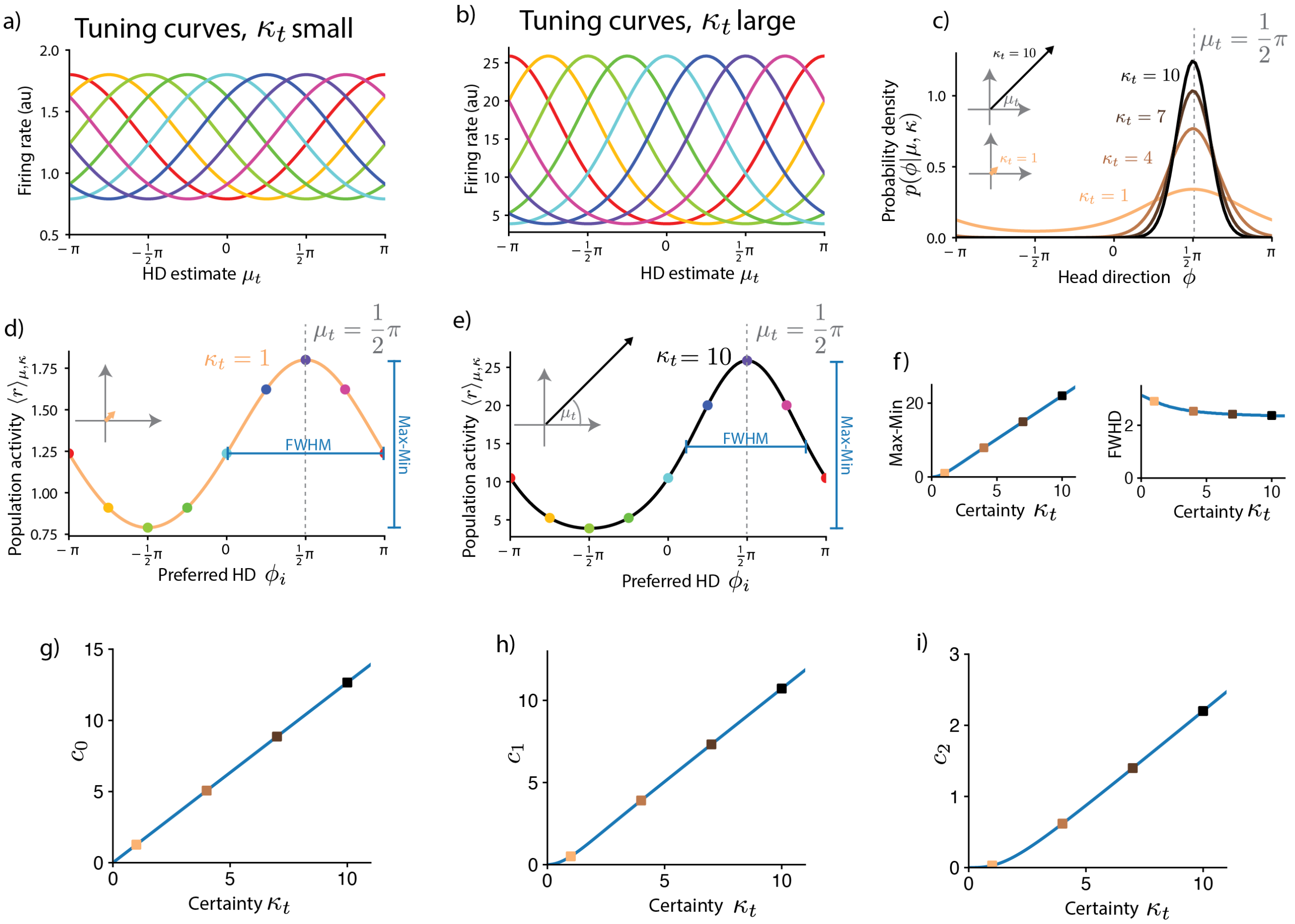

**Fig. S1. Encoding the HD with linear probabilistic population codes. a)** Tuning curves with respect to encoded HD estimate for small values of encoded certainty $\kappa_t$ are cosine-shaped. Here, we show tuning curves of 8 example neurons with $\kappa_t = 1$ (colors indicate preferred HD $\phi_i$). **b)** Tuning curves with respect to HD estimate for large values of encoded certainty $\kappa_t$ are von-Mises shaped (same 8 example neurons as in a, but for $\kappa_t = 10$). **c)** Von Mises probability densities for different values of encoded certainty $\kappa_t$ and fixed mean $\mu_t = \frac{\pi}{2}$. Note that the density sharpens around the mean with increasing certainty. Inset shows vector representation of a von Mises distribution with mean $\mu_t = \frac{\pi}{2}$, and, respectively, $\kappa_t = 10$ and $\kappa_t = 1$. **d)** Population activity profile (average neural firing rate conditioned on HD estimate $\mu_t$ and certainty $\kappa_t$) encoding the von Mises densities with mean $\mu_t = \frac{\pi}{2}$ and certainty $\kappa_t = 1$. Neurons are sorted by preferred HD $\phi_t$. Colored dots correspond to activity of neurons with tuning curves as in a). The phasor representation of the neural activity (inset) matches the vector representation of the encoded von Mises distribution in c. **e)** Population activity profile encoding the von Mises densities with mean $\mu_t = \frac{\pi}{2}$ and certainty $\kappa_t = 10$. **f)** Left: The amplitude (Max-Min) of the activity profile scales (approximately) linearly with certainty $\kappa_t$, except for very small values of $\kappa_t$. Right: The population activity bump's width (full width at half maximum, FWHM) is mostly unaffected by uncertainty $\kappa_t$, and saturates at a finite value for large $\kappa_t$, unlike the von Mises distribution it encodes (e.g., b), whose FWHM approaches zero for large values of $\kappa_t$. **g-j)** The Fourier component amplitudes of the population activity profile are mostly linear in encoded certainty $\kappa_t$, indicating that (i) the whole profile is scaled by $\kappa_t$, and that (ii) only focusing on the first Fourier component in our analysis is justified. For the tuning curves, we used $\xi = 1$ without loss of generality.

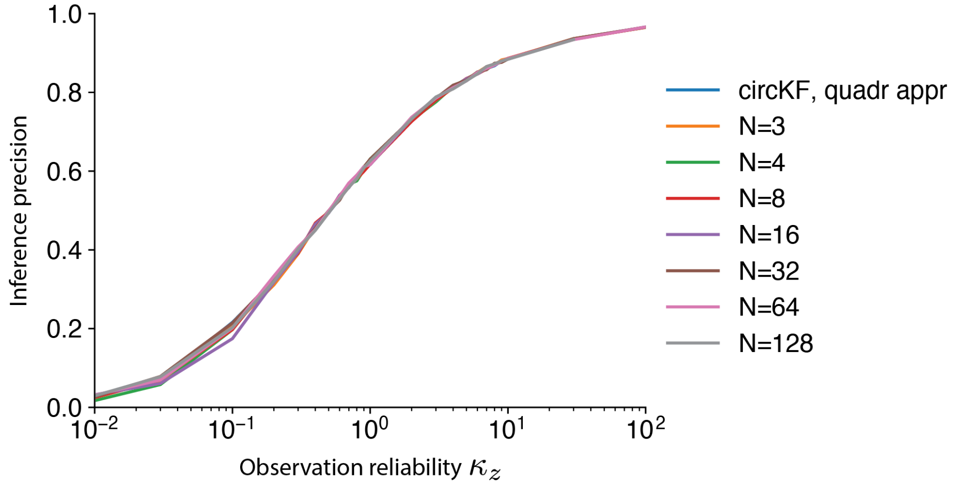

**Fig. S2. Network inference performance is mostly independent of the number of neurons $N$ in the Bayesian ring attractor network.** Here, for each value of the observation reliability $\kappa_z$ and number of neurons in the network $N$ we compute the circular average distance of the network's HD estimate $\mu_T$ from the true HD $\phi_T$ at the end of a simulation of length $T = 20$ from $P = 10'000$ simulated trajectories. The blue line (hidden below other lines) shows the performance of the quadratic approximation to the circular Kalman filter that the networks aim to implement. The network parameters of the single-population network in Eq. [S49] were those of the Bayesian ring attractor, i.e. $w_1^{\text{even}} = \frac{1}{\tau} + \frac{1}{\kappa_\phi + \kappa_v}$, $w_1^{\text{odd}} = \frac{\kappa_v}{\kappa_\phi + \kappa_v} v_t$, and $w^{\text{quad}} = \frac{1}{\kappa_\phi + \kappa_v}$. Other simulation parameters were: $\kappa_\phi = 1$, $\kappa_v = 1$, and $\Delta t = 0.01$.

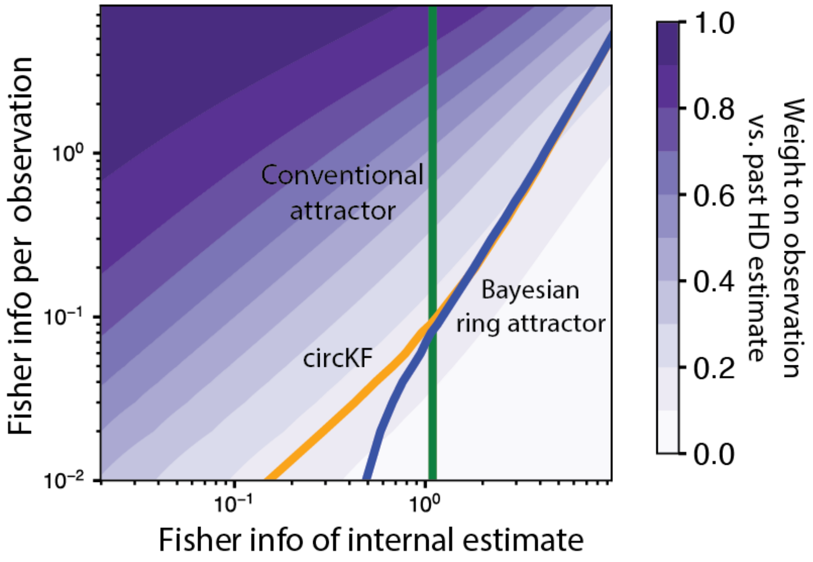

**Fig. S3. The weight with which a single observation contributes to the HD estimate varies with informativeness of both the HD observations and the current HD estimate.** Same plot as main text Fig. 4, only on a log-log scale. Here, we additionally plot the resulting updates for the circKF, to demonstrate that the Bayesian ring attractor (blue curve) only deviates from the circKF (yellow curve) for very uninformative observations.

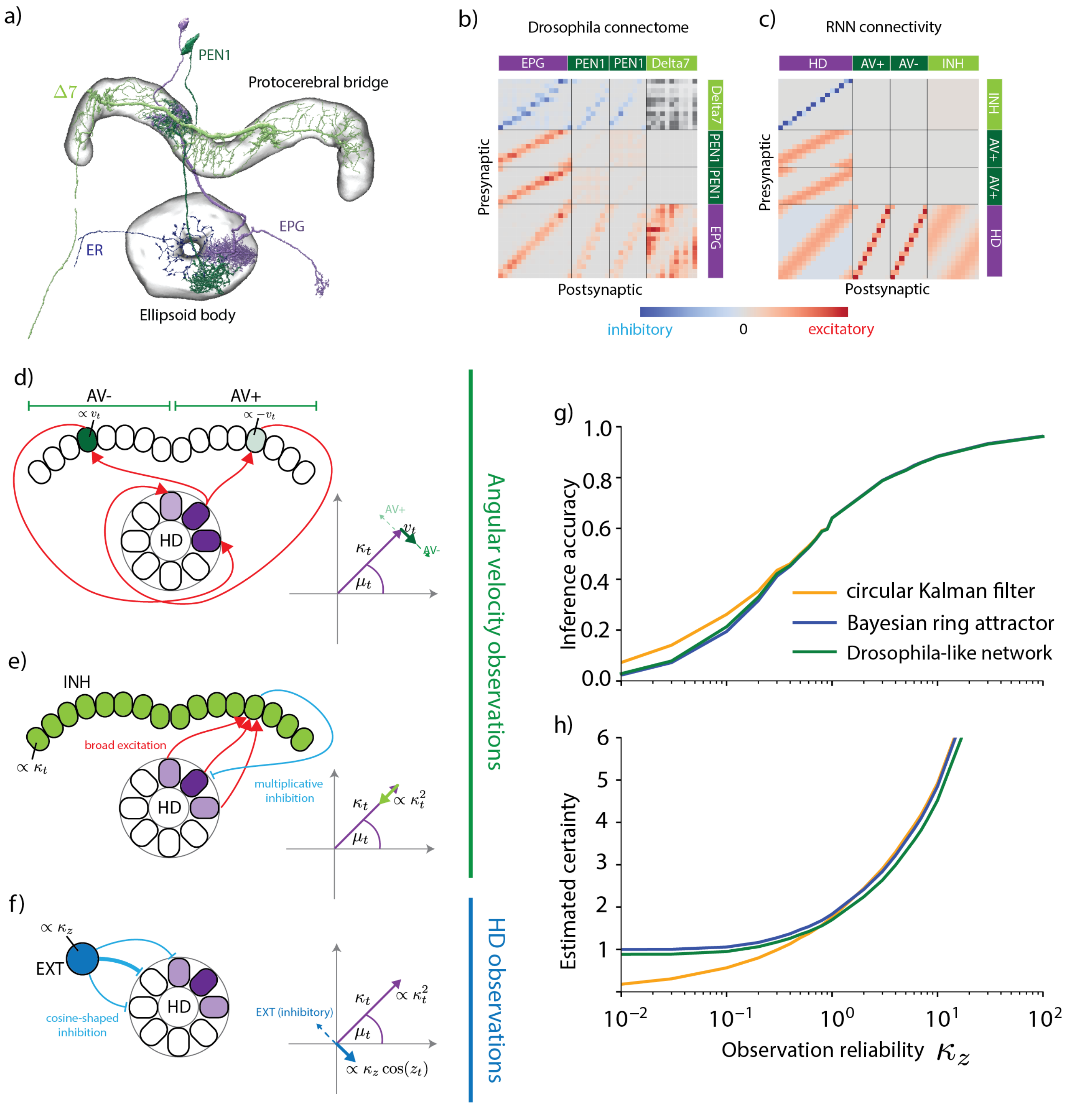

**Fig. S4. A Drosophila-like network implementing the circular Kalman filter. a)** Cell types in the Drosophila brain that could contribute to implementing the circular Kalman filter. **b)** Connectivity between EPG, $\Delta 7$ and PEN1 neurons, as recovered from the hemibrain:v1.2.1 database (10). ER neurons were omitted because they only form the inputs to the recurrently connected ring attractor. Here, neurons were grouped according to anatomical region as a proxy for preferred HD, and we used the total number of synaptic connections between two neurons to indicate connection strength. $\Delta 7$ to $\Delta 7$ connectivities are omitted, as the polarity of these connections (inhibitory or excitatory) remains unclear. **c)** The RNN connectivity profile that implements an approximation of Bayesian inference algorithm is strikingly similar to the connectivity of neurons in the Drosophila HD system. To avoid confusion with actual neurons, we refer to the neuronal populations in this idealized RNN as head direction (HD), angular velocity (AV+ and AV-, in reference to the two hemispheres), inhibitory (INH) and external input (EXT) populations. **d)** Differential activation of AV populations (left/right: high/low) in the two hemispheres as well as a shifted feedback connectivity from AV to HD populations effectively implement the odd (or shifted) connectivity needed to turn the bump position (here: clockwise shift for anti-clockwise turn). **e)** Broad excitation of the INH population by the HD population, together with a one-to-one multiplicative interaction between INH and HD population, implement the quadratic decay of the bump amplitude needed for the reduction in certainty arising from probabilistic path integration. **f)** External input is mediated by inhibiting HD neurons with preferred direction opposite the location of the HD observation, effectively implementing a vector sum of belief with HD observation. **g)** and **h)** The inference accuracy of the *Drosophila*-like network is indistinguishable from the Bayesian ring attractor. Here, we built a network from the connectivity profile shown in panel c, and used the following simulation parameters: $\kappa_v = 5$, $T = 20$, $\Delta t = 0.001$, results are averages over $P = 5'000$ simulations. Network architecture according to the full network in Eqs. [S67]-[S67], with baseline $o^{AV} = 0$, time scales $\tau_{HD} = 0.1$, $\tau_{AV+} = \tau_{AV-} = 0.01$, $\tau_{INH} = 0.001$, connection strengths $c_0^{HD} = -0.2$, $c_1^{HD} = 0$, $c^{AV^\pm \leftarrow HD} = 1$, $c_0^{INH \leftarrow HD} = 0.5$, $c_1^{INH \leftarrow HD} = -0.5$, $c_0^{INH} = 0.1$, $c_1^{INH} = 0$. Further, in the discretized dynamics we chose $N_{HD} = 100$, $N_{AV+} = 50$, $N_{AV-} = 50$, $N_{INH} = 100$, and $N_{EXT} = 100$.

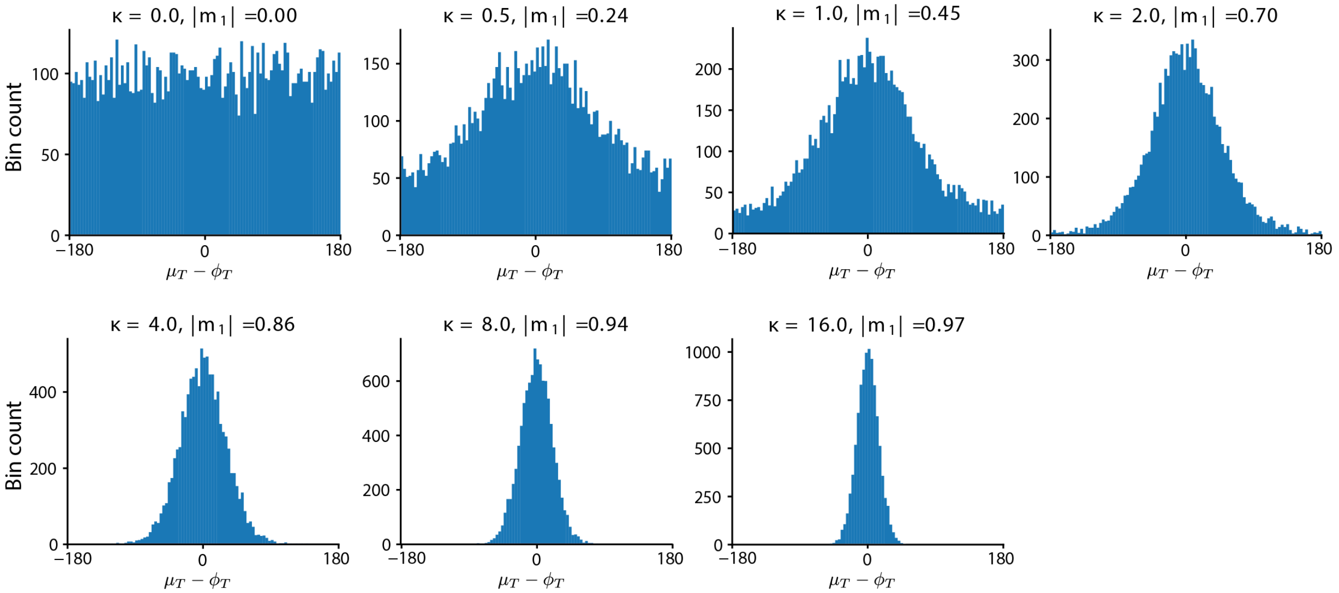

**Fig. S5. Visualizing the HD tracking perfomance measure.** To provide a better intuition for the used HD tracking performance measure we here show how a specific distribution of HD tracking errors (horizontal axis, in degrees) relates to this performance measure. In particular, we drew 10000 samples from a von Mises distribution $\mu_T - \phi_T \sim \mathcal{VM}(0, \kappa)$, where each drawn sample simulates one single deviation of the estimated HD (i.e., the mean of the filter posterior, $\mu_T$) from the actual, true HD, $\phi_T$. The different panels show the histogram of simulated errors for different $\kappa$'s (see panel headings). Our filtering performance measure, that is, the absolute value of the first circular average of the samples, can be computed for the von Mises distribution via $|m_1| = \frac{I_1(\kappa)}{I_0(\kappa)}$ (22). We confirmed numerically that this analytical expression matches the circular average empirically determined from these simulated errors. Simulating HD tracking errors by draws from a von Mises distribution was here only performed for convenience. The HD tracking errors arising in simulations of the filtering algorithms do not necessarily follow such a distribution.

#### References

1. Kutschireiter A, Rast L, Drugowitsch J (2022) Projection Filtering with Observed State Increments with Applications in Continuous-Time Circular Filtering. *IEEE Transactions on Signal Processing* pp. 1–1. Conference Name: IEEE Transactions on Signal Processing.
2. Brigo D, Hanzon B, Le Gland F (1999) Approximate nonlinear filtering by projection on exponential manifolds of densities. *Bernoulli* 5(3):495–534.
3. Gardiner CW (2009) *Stochastic methods: a handbook for the natural and social sciences*, Springer series in synergetics. (Springer, Berlin Heidelberg) No. 13, 4th ed edition.
4. Doucet A, Godsill S, Andrieu C (2010) On sequential Monte Carlo sampling methods for Bayesian filtering. *Statistics and Computing* p. 12.
5. Kutschireiter A, Surace SC, Pfister JP (2020) The Hitchhiker's guide to nonlinear filtering. *Journal of Mathematical Psychology* 94:102307.
6. Ma WJ, Beck JM, Latham PE, Pouget A (2006) Bayesian inference with probabilistic population codes. *Nature Neuroscience* 9(11):1432–8.
7. Beck JM, Latham PE, Pouget A (2011) Marginalization in Neural Circuits with Divisive Normalization. *Journal of Neuroscience* 31(43):15310–15319.
8. Pouget A, Beck JM, Ma WJ, Latham PE (2013) Probabilistic brains: knowns and unknowns. *Nature Neuroscience* 16(9):1170–8.
9. Dayan P, Abbott LF (2001) *Theoretical neuroscience: computational and mathematical modeling of neural systems*, Computational neuroscience. (Massachusetts Institute of Technology Press, Cambridge, Mass).
10. Scheffer LK, et al. (2020) A connectome and analysis of the adult Drosophila central brain. *eLife* 9:e57443. Publisher: eLife Sciences Publications, Ltd.
11. Hulse BK, et al. (2021) A connectome of the Drosophila central complex reveals network motifs suitable for flexible navigation and context-dependent action selection. *eLife* 10:e66039. Publisher: eLife Sciences Publications, Ltd.
12. Turner-Evans DB, et al. (2020) The Neuroanatomical Ultrastructure and Function of a Biological Ring Attractor. *Neuron* p. S0896627320306139.
13. Seelig JD, Jayaraman V (2015) Neural dynamics for landmark orientation and angular path integration. *Nature* 521(7551):186–191.
14. Turner-Evans D, et al. (2017) Angular velocity integration in a fly heading circuit. *eLife* 6:e23496.
15. Green J, et al. (2017) A neural circuit architecture for angular integration in Drosophila. *Nature* 546(7656):101–106. Publisher: Nature Publishing Group.
16. Omoto JJ, et al. (2017) Visual Input to the Drosophila Central Complex by Developmentally and Functionally Distinct Neuronal Populations. *Current Biology* 27(8):1098–1110.
17. Fisher YE, Lu J, D'Alessandro I, Wilson RI (2019) Sensorimotor experience remaps visual input to a heading-direction network. *Nature* (December 2018).
18. Kim SS, Hermundstad AM, Romani S, Abbott LF, Jayaraman V (2019) Generation of stable heading representations in diverse visual scenes. *Nature* (December 2018):1–6. Publisher: Springer US.
19. Hulse BK, Jayaraman V (2020) Mechanisms Underlying the Neural Computation of Head Direction. *Annual Review of Neuroscience* 43(1):31–54.
20. Skaggs W, Knierim J, Kudrimoti H, McNaughton B (1994) A model of the neural basis of the rats sense of direction in *Advances in neural information processing systems*, eds. Tesauro G, Touretzky D, Leen T. (MIT Press), Vol. 7.
21. Zhang K (1996) Representation of Spatial Orientation by the Intrinsic Dynamics of the Head-Direction Cell Ensemble: A Theory. *The Journal of Neuroscience* 16(6):2112–2126.
22. Mardia KV, Jupp PE (2000) *Directional Statistics*. (John Wiley & Sons). Pages: 3.
